# The combined impact of climate change, pesticides and pharmaceuticals on life history and behaviour in killifish (*Nothobranchius furzeri*)

**DOI:** 10.1101/2024.08.15.608117

**Authors:** Dongjin Kim

## Abstract

Animals in nature are confronted simultaneously with multiple stressors of both natural and anthropogenic origin including temperature stress, pollutants, predators and parasites. The overall goal of this study was to assess the sensitivity of fish to climate change in combination with both classic pollutants (pesticides) and emerging contaminants (pharmaceuticals). Specifically, we exposed individuals of the model fish species *Nothobranchius furzeri* to combinations of temperature fluctuation, 3,4-DCA and FLX. Our results indicate that both traditional endpoints such as body size and fecundity as well as behavioural endpoints are affected by daily temperature variation and by the tested pollutants. The fact that certain effects were much more prominent when pollutants were combined illustrates the relevance of stressor combination studies. Also, while the tested concentrations of pollutants did not cause excess mortality, they did induce significant behavioural changes which may carry serious fitness consequences. Ultimately, our study provides an effective illustration of the need for performing stressor combination studies and for including behavioural endpoints in ecotoxicological testing.

## 1. Introduction

### 1.1. Climate change

#### 1.1.1. Global warming

Various anthropogenic activities have led to atmospheric carbon dioxide (CO2) concentrations that are well above natural levels (Berger et al., 2012). Among other consequences, this has induced significant temperature increases in recent years since CO_2_, like other greenhouse gases, captures heat in the atmosphere (Berger et al., 2012). The global temperature has risen around 1°C since the Industrial Revolution (Figure 1.1). This temperature rise may impact organisms and as such threaten environmental integrity (McMichael et al., 2003).

**Figure 1.1.**
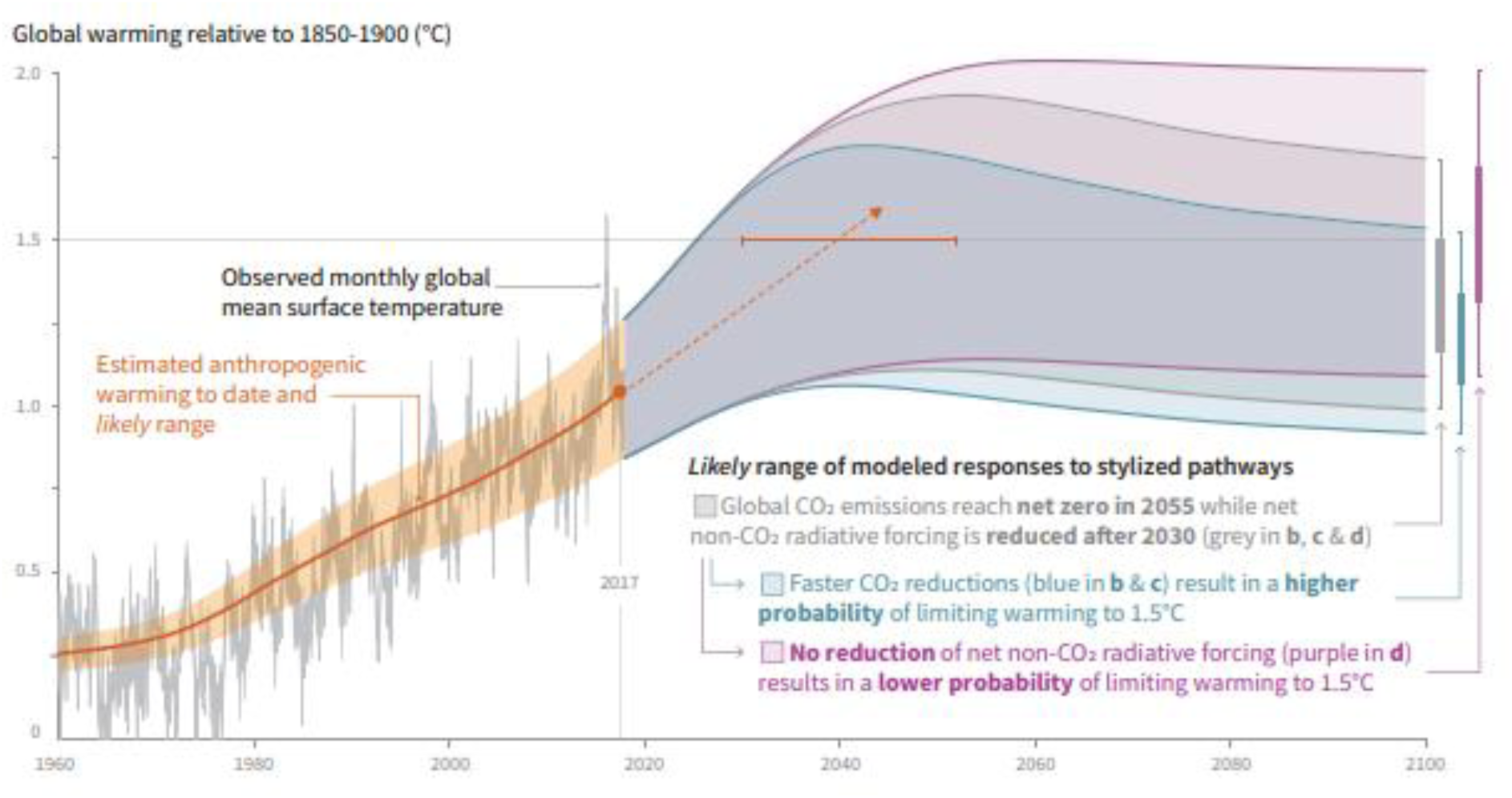
A model of global warming based on anthropogenic emission (Masson-Delmotte et al., 2018)

The Intergovernmental Panel on Climate Change (IPCC) has set a target to limit temperature rise to 1.5°C compared to the Industrial Revolution to minimise unfavourable consequences of climate change (Masson-Delmotte et al., 2018). IPCC has developed several scenarios of global warming until 2100 depending on different levels of carbon emission in the atmosphere as shown in Figure 1.1. Most of the projections forecast temperature rise, the magnitude of which depends on the specific adopted scenario.

#### 1.1.2. Overall impacts on ecosystem

Climate change is expected to impact ecosystems around the globe (Cáceres, 2012). In addition to the associated temperature stress, organisms may have to cope with changing precipitation patterns and disrupted seasonal cues in the face of climate change (Windisch et al. 2014). Moreover, climate change may also drive new indirect stressors, including higher levels of inter- and intraspecific competition for depleting resources and the spread of infectious diseases (Cáceres, 2012).

The unfavourable effects of climate changes could induce extinctions and geographical displacements on a massive scale (Dawson et al., 2011). Hence humans and animals are obliged to adapt to new challenges driven by climate change including depletion of resources, food chain disruption and transition in pathogen-host interaction (Cáceres, 2012; McMichael et al., 2003). Consequently, biodiversity could be threatened and proper ecosystem functioning compromised. (Dawson et al., 2011).

#### 1.1.3. Pollution

Global anthropogenic change does not only entail temperature change but also, for instance, increasing chemical pollution (Baste et al., 2012). Pollution is one of the main drivers of global change and a key environmental problem. When it happens, a plethora of chemical products is released in the environment, where they may affect organisms and threaten the integrity of the ecosystem (Baste et al., 2012).

In freshwater systems, the impact of pollutants including pesticides and heavy metals continues to increase (Brown et al., 2015). In addition, pharmaceutical products such as endocrine-disrupting chemicals are often reported in surface waters (Birnbaum, 2013).

### 1.2. Ecotoxicology

#### 1.2.1. Research area and methods

Ecotoxicology has been derived from the combination of ecology and toxicology to improve our understanding the negative effects of chemical compounds in an environment, in an overall attempt to prevent environmental degradation through science-based policy (Walker et al., 2012). In ecotoxicological studies, the impact of chemical compounds on the ecosystem is assessed by performing standardised tests across multiple levels of biological organisations; molecular, cellular, organismal, population and community (Fend, 2001).

To this end, model organisms are exposed for a certain amount of time to series of dilutions of a certain compound to examine toxicity and environmental risk (Holmstrup et al., 2010). Among several established tests, the Organisation for Economic Co-operation and Development (OECD) has developed the fish acute toxicity test (OECD Test Guideline No. 301) (Selck et al. 2002) which are designed to facilitate result-comparison across studies and provide rapid ‘ready-to-use’ toxicity data for regulatory decision-making.

Typically, several parameters of compound toxicity can be obtained from these tests by scoring life history responses and dose-response curves (Figure 1.2). The lowest-observed-effect concentration (LOEC) is the lowest experimentally tested concentration at which an observable effect can be detected (Selck et al., 2002). The concentration of the compound at which 50% of the tested organisms die, is referred to as the LC50-value (‘Lethal Concentration’) (Ferrari et al., 2003). When any other effect but mortality is assessed (e.g. physiological endpoints, behavioural endpoints), this value is more generally referred to as the EC50-value (‘Effect Concentration’) (Ferrari et al., 2003).

**Figure 1.2.**
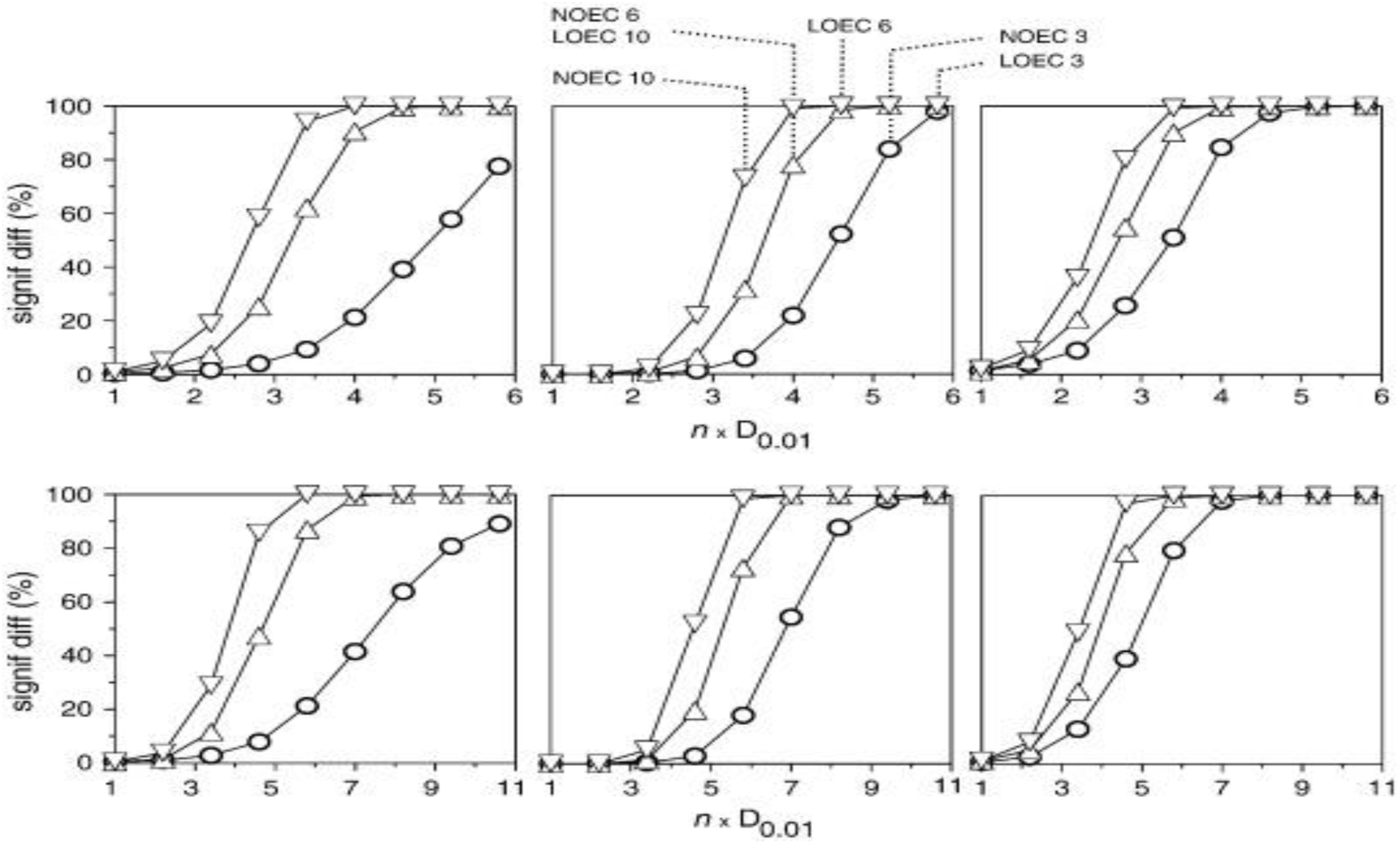
Dose-response curves with lowest (LOEC) and no observed effect concentration (NOEC) (Murado & Prieto, 2013)

#### 1.2.2. Shortcomings of traditional ecotoxicological testing

Standard tests in aquatic ecotoxicology are aimed at predicting the risk of chemical compounds in the aquatic environment (Brodin et al., 2013). However, the scope of these tests is usually tied up in the assessment of mortality and stress on animal species and other responses like behavioural alterations are typically neglected (Holmstrup et al., 2010). Yet also these are highly relevant since they may result in severe fitness effects (Klaminder et al., 2014).

Furthermore, unlike laboratory environments, organisms usually encounter sub-optimal conditions in their natural environment (Chapman et al., 1998). Therefore, chemical compounds sometimes affect organisms much more severely in nature (Chapman et al., 1998).

Organisms in nature are usually also exposed to stressors for longer periods than in laboratory tests which often only run for a couple of days. There is a need for whole-life and multi-generational testing the validation of safety margins for compounds and to assess effects of acclimatisation, habituation, adaptation but also transgenerational effects (Holmstrup et al., 2010). Finally, while in classic ecotoxicology temperature is usually kept stable, especially shallow waters typically experience strong daily and seasonal temperature fluctuations and the relevance of this variation in relation to estimating compound effects should be investigated (Klaminder et al., 2014).

### 1.3. Contaminants

#### 1.3.1. Pesticides as traditional contaminants

Pesticides are used worldwide in agriculture to increase crop productivity (Kattwinkel et al., 2011). Three types of pesticides – fungicides, herbicides and insecticides – can be applied by farmers depending on the purpose and characteristics of crops (Figure 1.3; Carvalho, 2017). They enter aquatic systems such as streams and rivers through run-off (Carvalho, 2017) industrial spilling, direct application or penetration. (Schriever & Liess, 2007). Pesticides are expected to increasingly end up in the environment and their effects may be exacerbated by temperature rise as expected under climate change (Kattwinkel et al., 2011).

**Figure 1.3.**
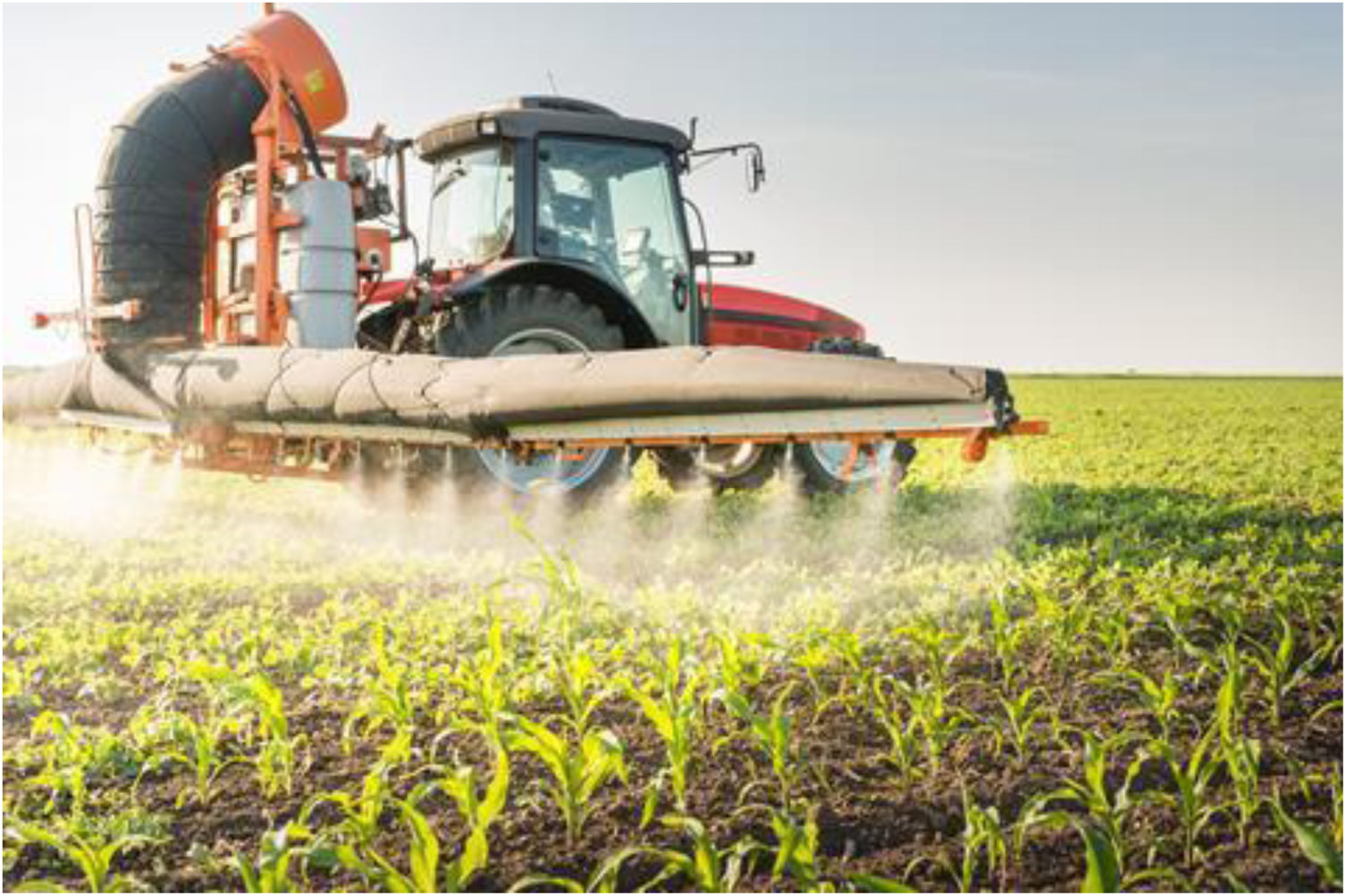
Use of pesticides (Burke, 2017)

Pesticides not only impact target species, but also affect non-target species when they end up in the natural environment. (Bretaud et al., 2000). As intense precipitation events are expected to increase in certain regions (Masson-Delmotte et al., 2018), the residue of pesticides in eroded sediments and run-off water is expected to be more often transported to soil and aquatic environments (Noyes et al., 2009).

#### 1.3.2. 3,4-dichloroaniline

3,4-dichloroaniline (3,4-DCA) is a precursor or a by-product in the production of herbicides that is commonly used on a worldwide scale. Its molecular formula is C_6_H_5_Cl_2_N (Figure 1.4). The compound is for instance frequently used in rice cultivation to control the growth of weeds, especially in Asia and North America (Xiao et al., 2016). The residue of 3,4-DCA had been found up to a depth of 30 cm depth in soil from rice paddies (Kearney et al., 1970).

**Figure 1.4.**
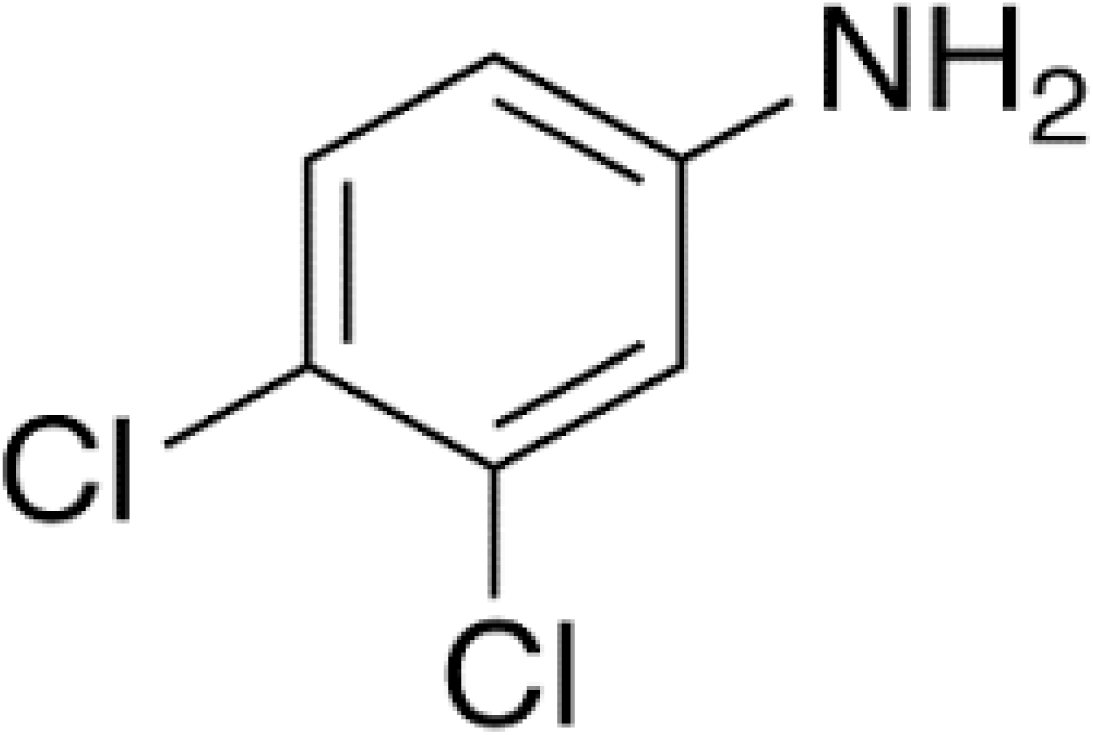
The chemical structure of 3,4-dichloroaniline (TRC, 2020)

3,4-DCA has a low solubility in water and can be converted to 3,4,3ʹ,4ʹtetrachloroazobenzene (TCAB) via microbial peroxidases. TCAB is known to have detrimental impacts on aquatic organisms (Allinson & Morita, 1995a & 1995b) including aquatic snails (*Indoplanorbis exustus*) and Japanese Medaka (*Oryzia latipes*). Known effects included a reduced food intake and body size.

#### 1.3.3. Pharmaceuticals as emerging contaminants

Pharmaceuticals are biologically active chemicals that are aimed at alleviating symptoms of disease and treating diseases (Healthline, 2020). Due to their increasing and continuous use, pharmaceutical compounds ever more frequently end up in the environment (Kümmerer, 2010).

Pharmaceuticals are generally resistant to biodegradation and excreted by humans without or with very little chemical degradation compared to the original compounds (Arnold et al., 2014) as shown in Figure 1.5 (Reid et al., 2005). Many pharmaceuticals are also used to increase livestock or agricultural productivity across the globe (Boxall, 2009). As drug target molecules are often evolutionary conserved across the animal kingdom, non-target species in the environment are likely to be affected by pharmaceutical pollution through specific biological effects (Gunnarsson et al., 2008).

**Figure 1.5.**
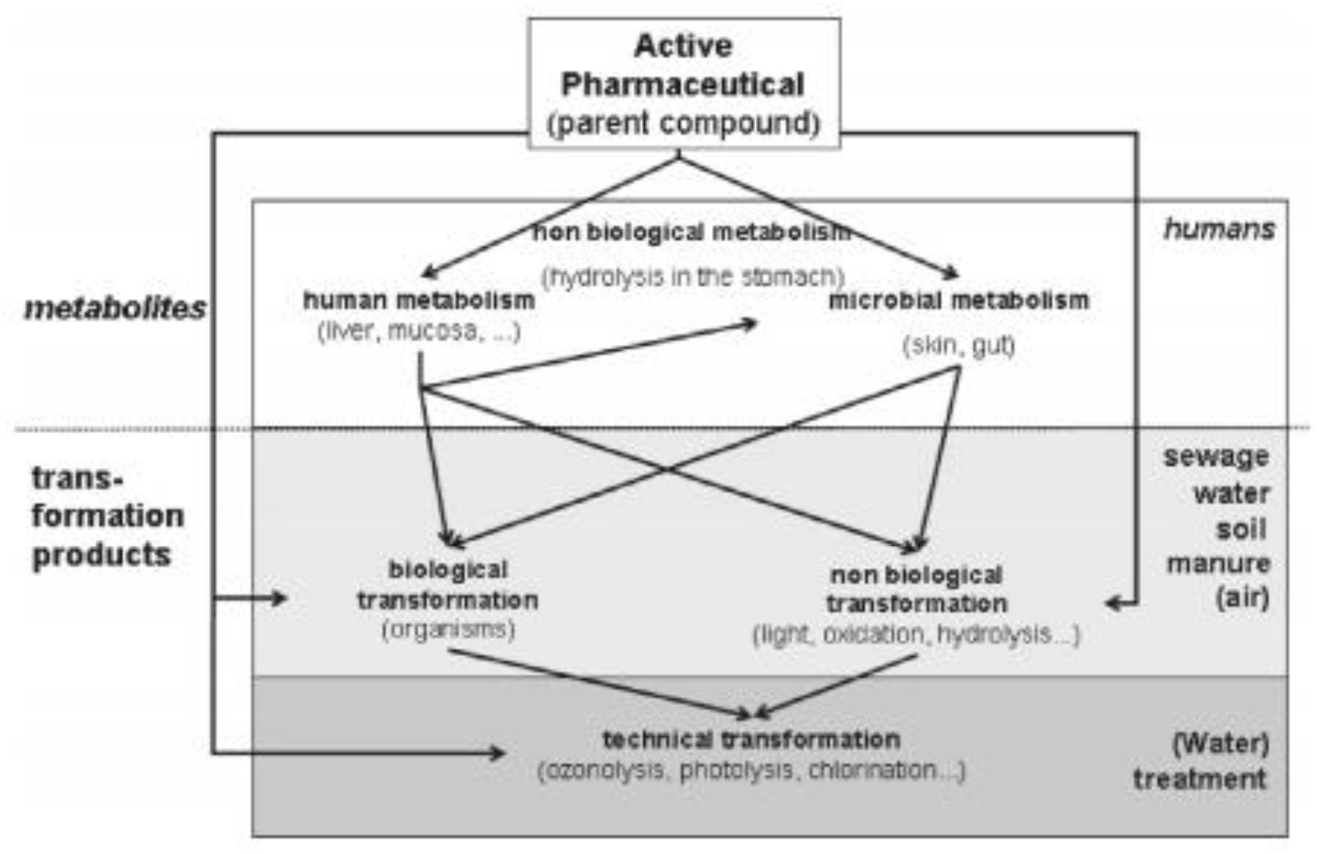
Pathways of pharmaceutical product transformation and degradations (Kümmerer, 2010)

#### 1.3.4. Fluoxetine

Serotonin is a neurotransmitter that regulates many physiological processes and is known to strongly influence emotions (Healthline, 2020). Fluoxetine (FLX) is a pharmaceutical compound with a molecular formula of C_17_H_18_F_3_NO (Figure 1.6) that has antidepressant effects and functions as selective serotonin reuptake inhibitor (SSRI) (Healthline, 2020). It prevents the reuptake of the neurotransmitter serotonin in the synaptic cleft, leading to higher levels of extracellular serotonin (Malagié et al., 1995; Altamura et al., 1994). A dose of FLX reduces an uptake of nutrition and alleviates anxiousness.

**Figure 1.6.**
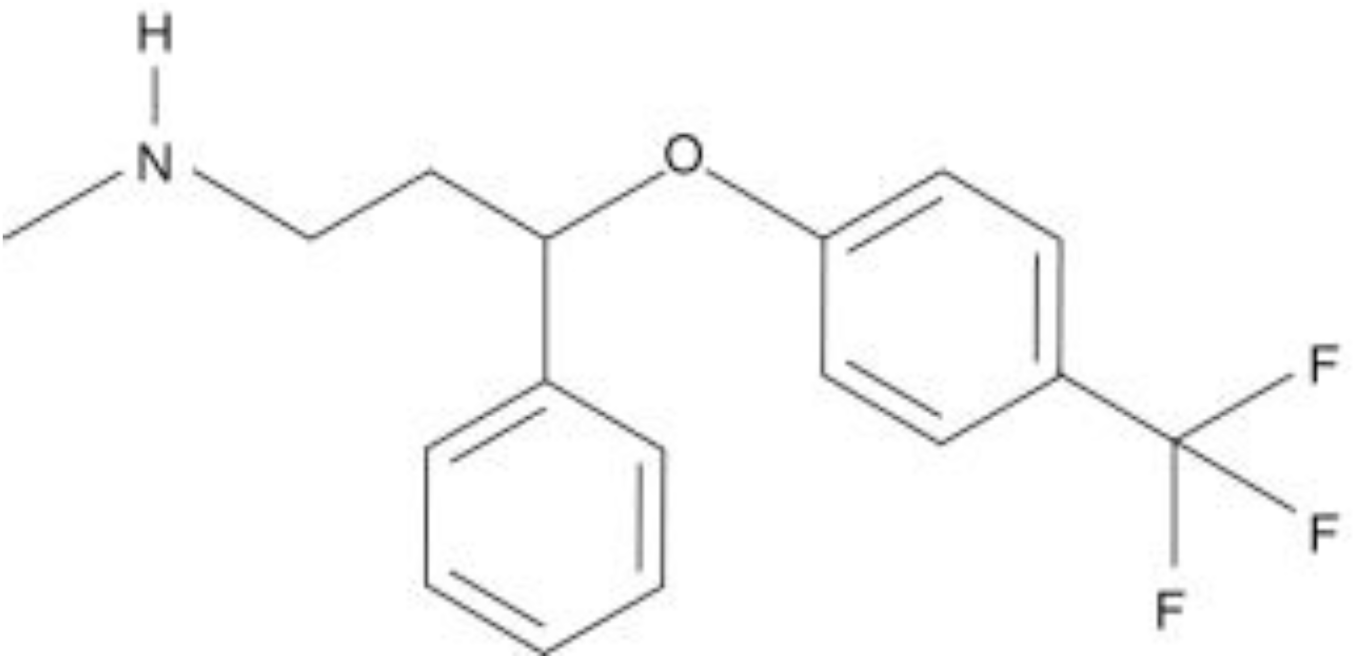
The chemical structure of FLX (Waldman, 2009)

Increasing concentrations of FLX are found in nature (Brodin et al., 2014). FLX is metabolised in the human body and excreted during urination while the composition remains practically unchanged (Puckowski et al., 2016). In surface waters, FLX concentrations are expected to further increase because of increasing human use and inadequate removal of pharmaceutical residues from sewage (Reid et al., 2005; Kümmerer, 2010).

### 1.4. Animal behaviour in ecotoxicology

Behaviour can be considered an integrative trait and comprises observable actions and changes of an organism, regulated by genes, neurons and anatomy in response to the environment (Figure 1.7; Gomez-Marin et al., 2014).

**Figure 1.7.**
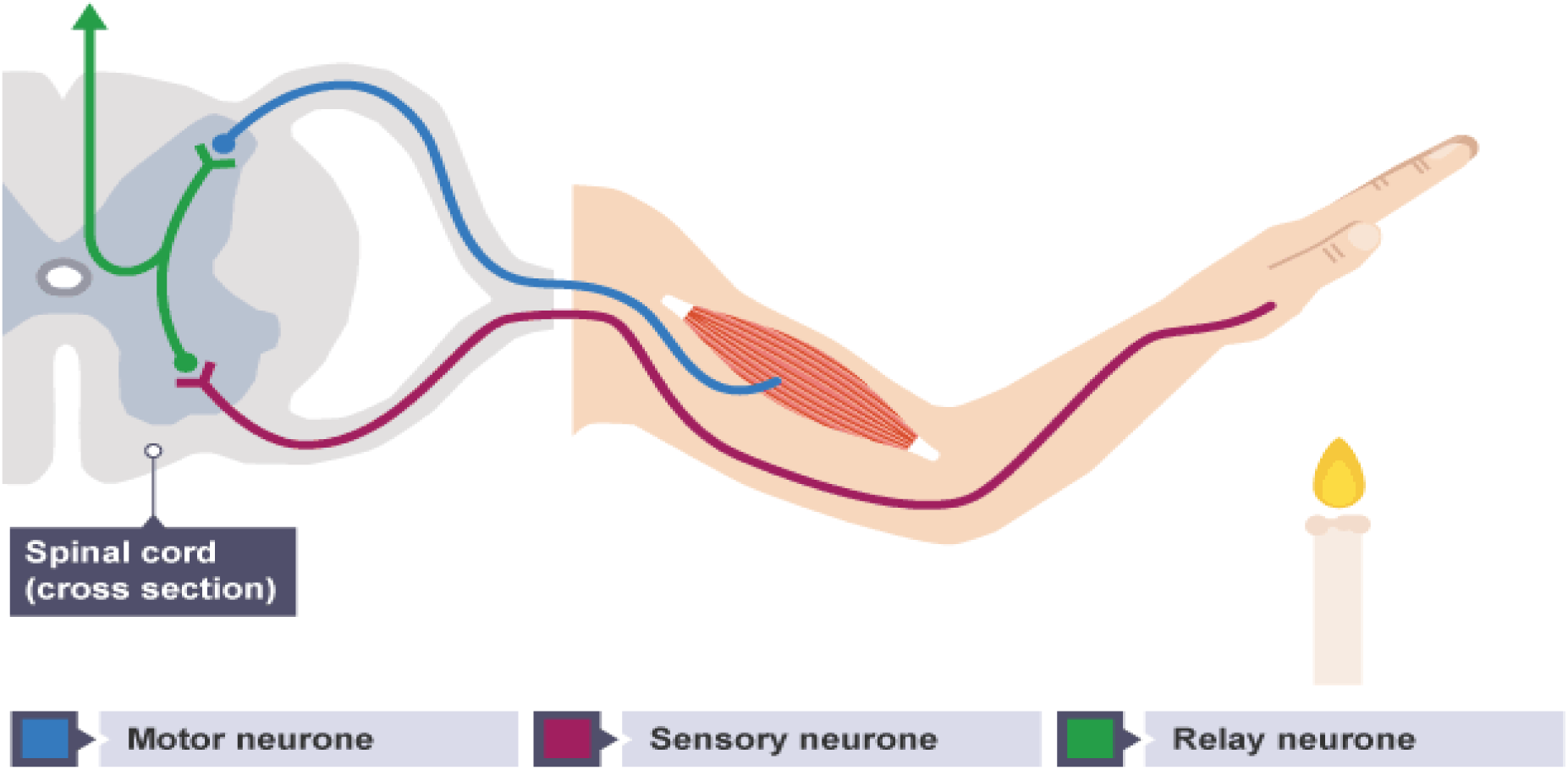
A stimulus is detected by receptors and transferred through neurons after which an effector determines a response of behaviour. Behaviour is the response that is externally shown (BBC, 2017).

Behaviour is tightly linked to the performance and fitness of organisms (Tai et al., 2012). Behaviour can be altered to adapt to diverse conditions including natural or anthropogenic environmental triggers (Gomez-Marin et al., 2014). Combined effects of global warming and contaminants could cause negative effects on organisms and impact their behaviour. (Klaminder et al., 2014; Gomez-Marin et al., 2014).

Behavioural traits – aggressiveness, boldness and sociality – have been reported to be significantly impacted by exposure to pharmaceuticals in both low and high concentrations (Brodin et al., 2013). For example, Arabian killifish (*Aphanius dispar*) exposed to 3µg/L of FLX were less active compared to unexposed individuals. (Barry, 2013).

### 1.5. Nothobranchius furzeri

#### 1.5.1. Characteristics

*Nothobranchius furzeri* (Turquoise killifish) is an annual fish that inhabits temporary pond systems across South-eastern Africa (mainly Mozambique and Zimbabwe; Figure 1.8). *N. furzeri* occurs in the area between the Save river and Lebombo ridge (Reichard et al., 2009). The average annual precipitation of this region varies from 200 mm to 600 mm (Tozzini et al., 2013). Temporary ponds in the region typically form after heavy rainfall events but they are short-lived due to the high temperatures and tend to dry in weeks (Cellerino et al., 2015).

**Figure 1.8.**
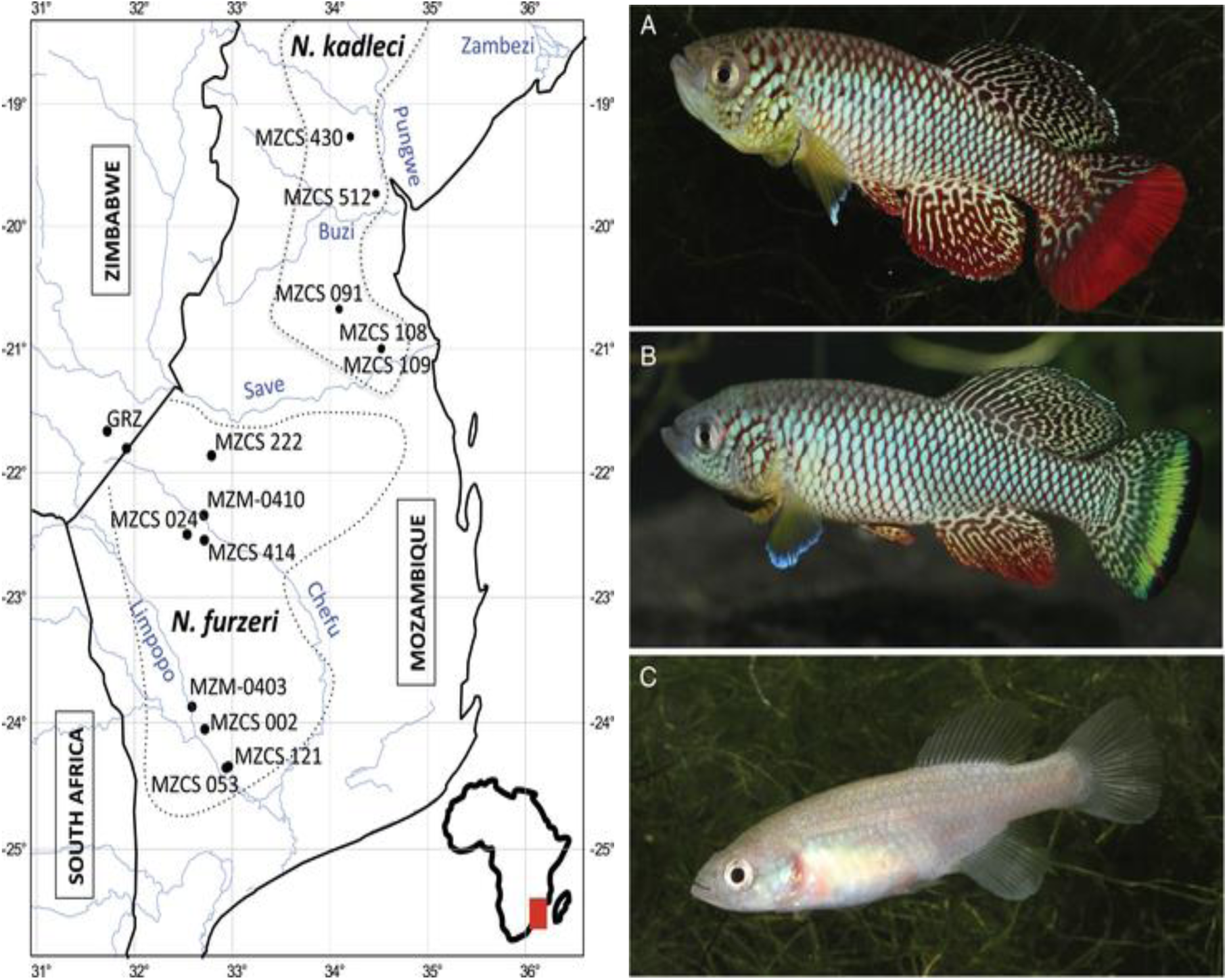
<left> Distribution of *N. Furzeri* (Cellerino et al., 2016) <right> Adult male with red tail (A), yellow tail (B) and female (C) of *N. Furzeri* (Cellerino et al., 2016).

*N. furzeri* is among the shortest-lived vertebrates, maturing in 3 weeks and dying in less than 12 months (Polačik & Reichard, 2011). *N. furzeri* has a small body size of about 3 – 15 cm, with mature males being both larger and brighter than females. There are two male colour morphs – a morph characterised by a red caudal fin and a morph yellow caudal fin (Figure 1.8; Cellerino et al., 2016). The bright colouration of males likely relates to sexual selection as this can attract females by improving visibility in the turbid pond environment (Cellerino et al., 2016).

At the onset of the rainy season and when pools are inundated, eggs immediately hatch. Hatchlings rapidly grow and reach maturity within 3 weeks after hatching (Haas, 1976). Mature individuals spawn daily and produce drought-resistant eggs before the end of the rainy season and before their pools dry up again (Cellerino et al., 2016). Eggs can remain dormant in the sediment until a next inundation event (Figure 1.9; Cellerino et al., 2016).

**Figure 1.9.**
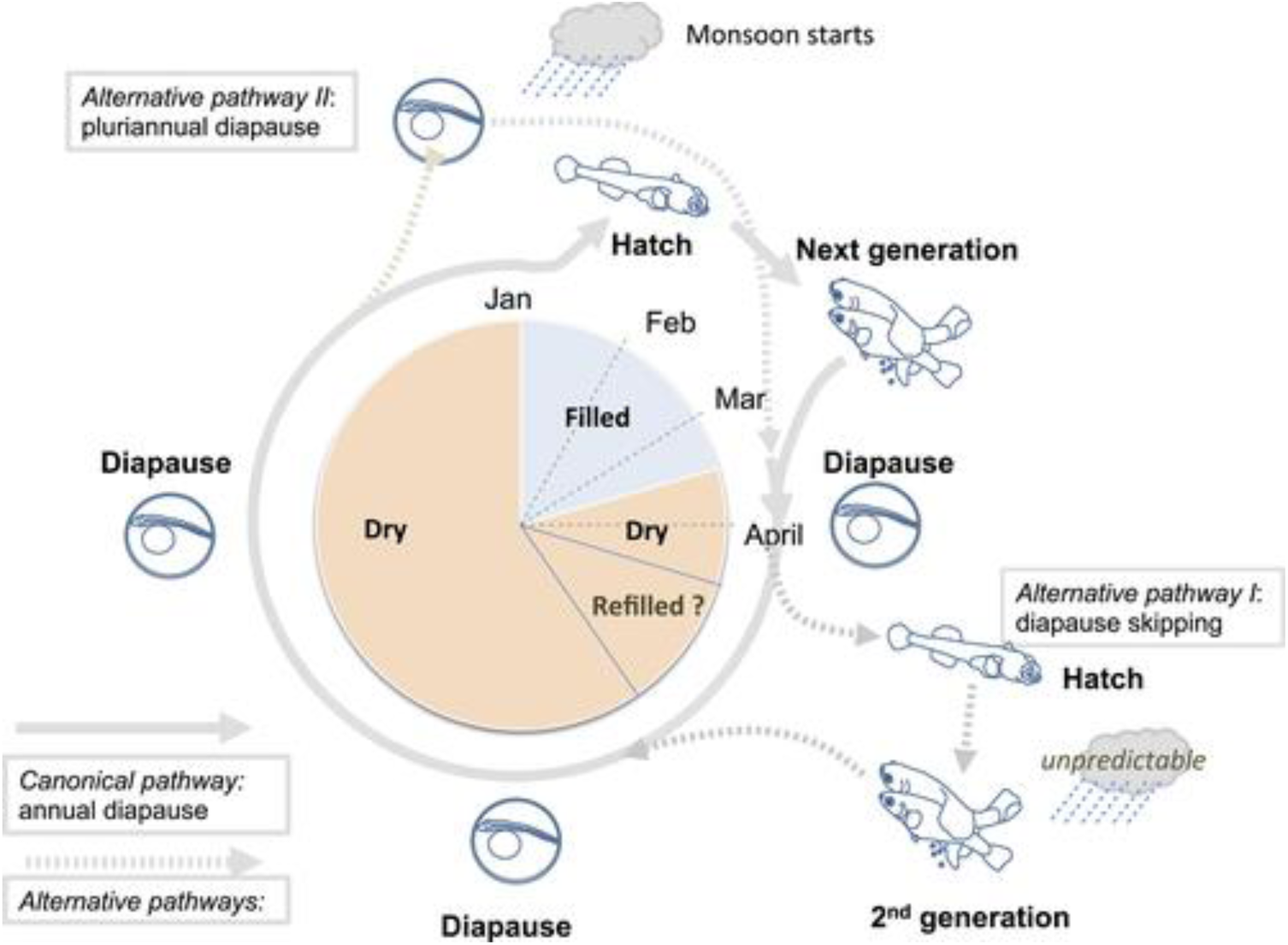
A life cycle of *N. furzeri* (Cellerino et al., 2016)

#### 1.5.2. Model species

*N. furzeri* has become a popular model species for biological research due to its short lifespan and fast maturation (Cellerino et al., 2016). Culturing the species in the laboratory comes with numerous advantages of which fast maturation and short lifespan are the most important ones since this enables life history and multi-generational studies in a time-efficient way. Next to that, since the fish produce long lived eggs, there is no need for continuous large cultures which reduces laboratory animal use and costly culture maintenance (Cellerino et al., 2016).

The short lifespan of *N. furzeri* allows examination of effects of contaminants of chronic exposure and even transgenerational effects and effects of exposure across consecutive generations in a relatively short lifespan. Over the past years, *N. furzeri* has been introduced as a new model in ecotoxicological testing (Philippe et al., 2018) and is fast gaining popularity as a model species in the field of behavioural ecotoxicology (Thoré et al., 2018; Thoré et al., 2019; Cellerino et al., 2016).

### 1.6. Goals and hypothesis

The goal of this study is to examine the separate and combined behavioural and life-history impact of whole-life mixed-stressor exposure to a conventional contaminant (the pesticide 3,4-DCA) and an emerging contaminant (the pharmaceutical fluoxetine) in the turquoise killifish (*N. furzeri*) under different temperature regimes. In addition, we aim to evaluate the potential of *N. furzeri* as a model for whole-life and multigenerational mixed-stressor exposure studies in a climate change context. As organisms are seldom exposed to single stressors in the environment, we here explore the impact of combined exposure to 3,4-DCA, fluoxetine and different temperature regimes (constant and daily varying). Overall, we expect both compounds to impact life-history and behavioural expression as well as hypothesise the emergence of, larger than additive, effects between the compounds.

Given that many drug target molecules are evolutionary conserved among vertebrate taxa (Gunnarsson et al. 2016), and considering the anxiolytic and anorexigenic properties of fluoxetine, we expect that fluoxetine exposure will induce effects in *N. furzeri*. Specifically, we expect an increase in reproductive output upon exposure, as recently demonstrated in a range of other fish species. For instance, Martin et al. (2019) illustrated that FLX exposure resulted in higher sperm quality and mating behaviour of mosquitofish (*Gabusia holbrooki*). We furthermore hypothesise a smaller growth rate underpinned by a reduction in feeding in fluoxetine-exposed fish. This hypothesis is supported by Thoré et al. (2019), who showed that feeding-behaviour in *N. furzeri* was impaired after short-term FLX exposure. Furthermore, we expect that long-term exposure to the compound will result in a decreased growth rate. This has been shown for instance by Hazelton et al. (2014) who found that FLX exposed mussels (Lampsilis fasciola) had a smaller body size than the control group individuals.

Since 3,4-DCA mainly causes tissue hypoxia as it reduces the oxygen binding affinity of haemoglobin (Allinson & Morita, 1995a; Allinson & Morita 1995b), we expect that 3,4-DCA exposure will induce negative effects in *N. furzeri*. For instance, we expect a decrease in reproductive output upon exposure. According to Preuss et al. (2010), exposure of *D. magna* to 3,4-DCA resulted in reduced reproductive output and more aborted eggs. We also expect that feeding and locomotor activity will be impaired as was shown by Velki et al. (2017) for zebrafish (*Danio rerio*) after exposure to 3,4-DCA.

Temperature fluctuation forces ectothermic organisms to continuously re-adapt their physiology to the surrounding conditions (Cáceres, 2012). We run the experiment under both constant and daily varying temperature conditions. Through combining the temperature regimes with both 3,4-DCA and FLX in a full factorial design, we can assess potential interactive effects of temperature regimes with these pollutants and among the pollutants themselves. In addition, by also including control conditions in absence of pollutants we can directly assess temperature effects. We expect that behaviour will be altered by exposure to the contaminants and that this impact will be exacerbated under fluctuating temperature conditions.

## 2. Material and methods

### 2.1. Breeding *N. furzeri*

The experiment ran from 18 November 2019 to 20 March 2020. At the onset of the experiment, 80 ready-to-hatch eggs of the GRZ-AD laboratory line reared at Animal Museum, KU Leuven were inundated in a 2L hatching tank at 26°C, following the protocol by Philippe et al. (2019). We used aerated reconstituted water at a 600 ± 10 µS/cm conductivity (type II RO water mixed with Instant Ocean Salt) enriched with 1 g/L humic acid as a hatching medium.

8 days post-hatching, hatchlings were individually transferred to a 2L transparent glass jar to facilitate individual monitoring (Figure 2.2). Throughout the experiment, fish were kept in temperature-controlled incubators at a light: dark regime of 14: 10 hours.

**Figure 2.1.**
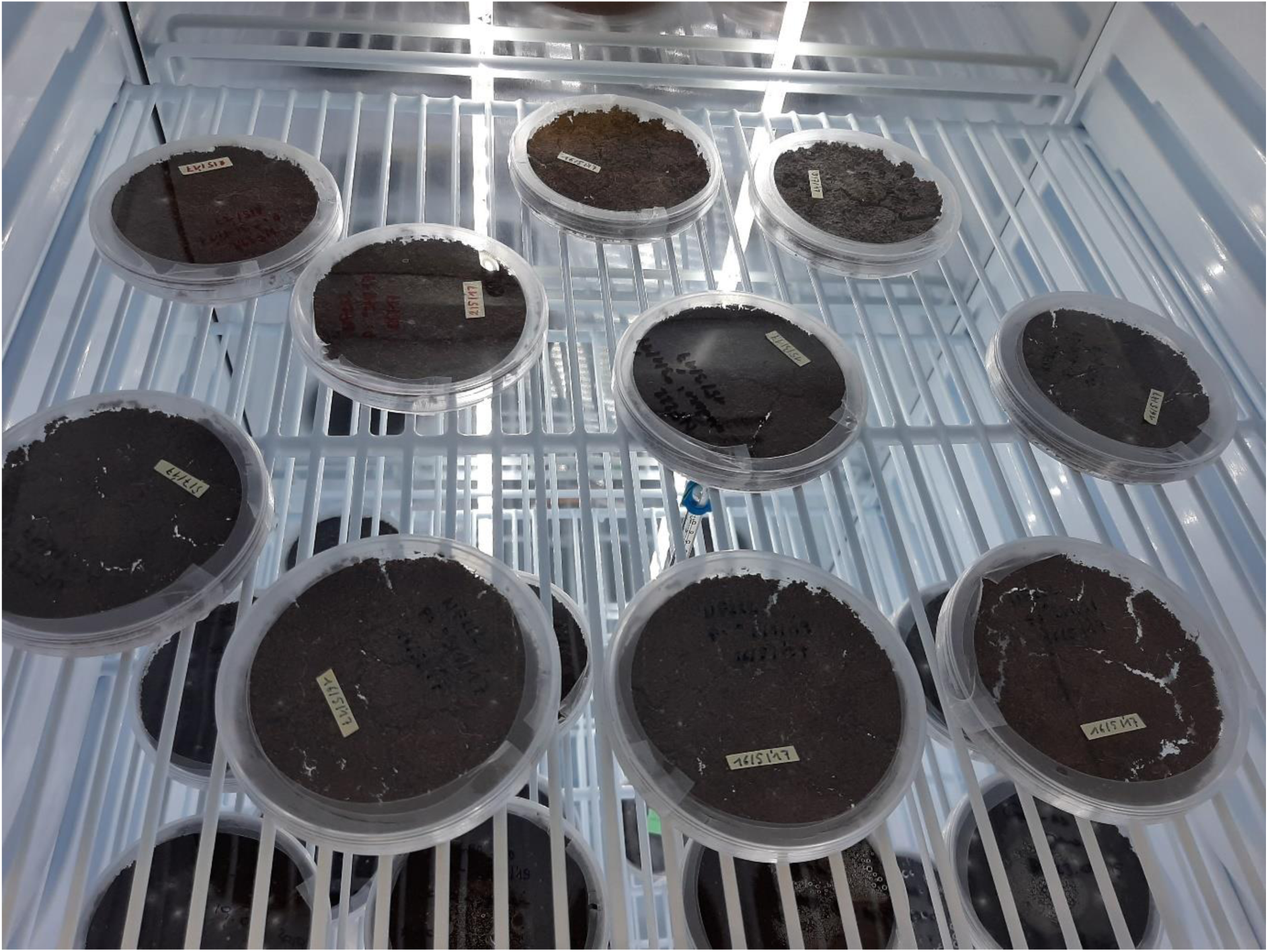
Eggs stored in an incubator at 28°C and ready to be hatched.

**Figure 2.2.**
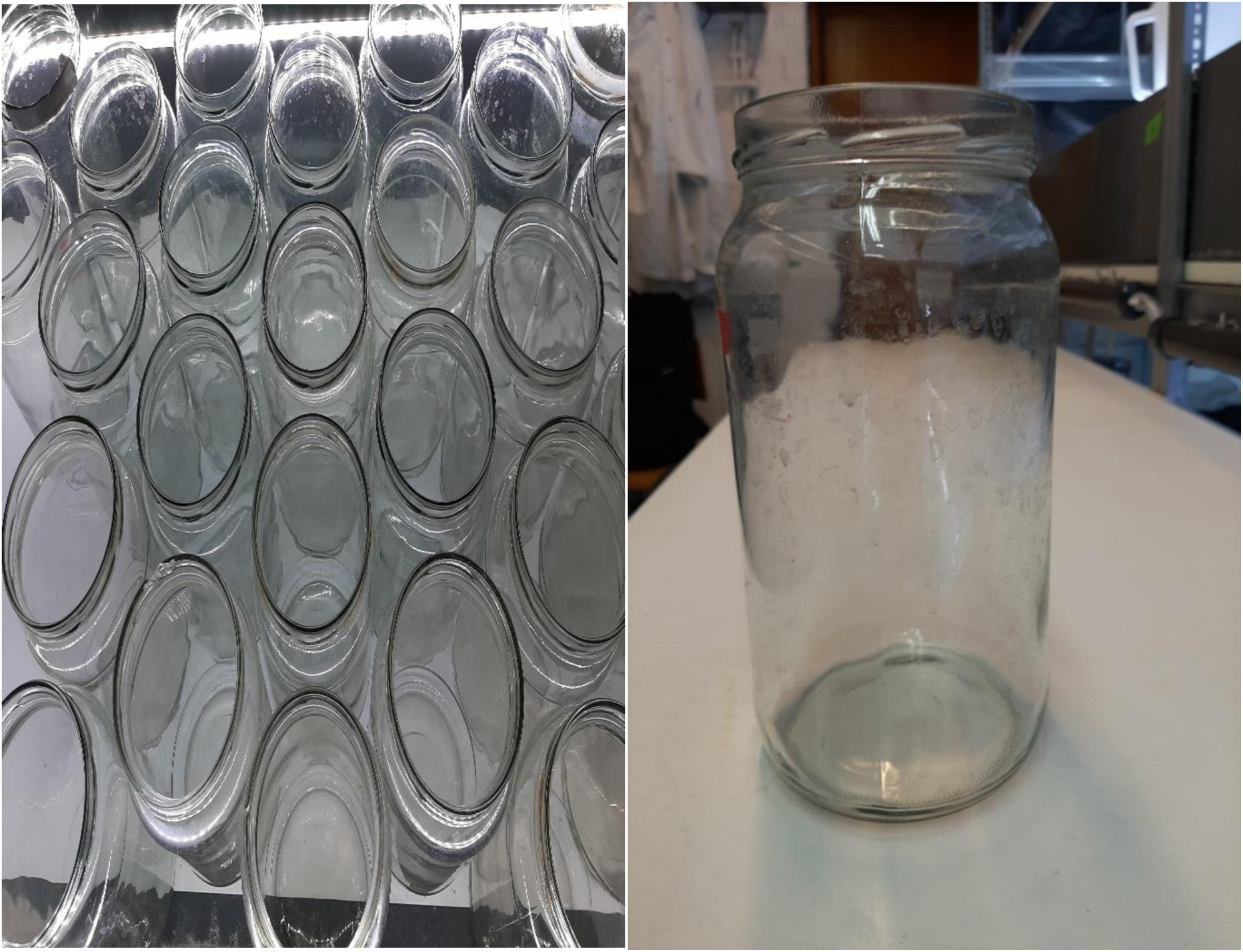
2L jars for storage of individual fish

Fish were fed twice daily an ad libitum quantity of *Artemia franciscana* nauplii (Ocean Nutrition, Essen, Belgium) from Day 2 to Day 20. Fish were subsequently weaned on *Chironomus* larvae (Ocean Nutrition, Essen, Belgium) and *Artemia franciscana* nauplii for a week, and exclusively fed an ad libitum quantity of *Chironomus* larvae as of Day 21 until the end of the experiment. Water was refreshed twice in a week throughout the experiment.

### 2.2. Experimental setup

Individuals were randomly subdivided across two temperature regimes; a daily fluctuating (DTV of 23-30°C) and a constant temperature (26°C). Both groups were kept at a 14 hour light: 10 hour dark regime to match the natural environment in the summer of South-eastern Africa (Table 2.1). Temperatures in the variable temperature regime were logged using data loggers (ONSET 1-800-loggers, 2020) to validate the real temperatures.

**Table 2.1.** Temperature and daylight setting for the group of temperature fluctuation.

| <u>Time</u> | <u>Temperature</u> | <u>Diurnal-nocturnal cycle</u> |
| --- | --- | --- |
| 4:00 | 23 | Darkness |
| 5:00 | 23.5 | Light |
| 6:00 | 24 | Light |
| 7:00 | 25 | Light |
| 8:00 | 26 | Light |
| 9:00 | 27 | Light |
| 10:00 | 28 | Light |
| 11:00 | 29 | Light |
| 12:00 | 30 | Light |
| 13:00 | 30 | Light |
| 14:00 | 30 | Light |
| 15:00 | 29.5 | Light |
| 16:00 | 29 | Light |
| 17:00 | 28.5 | Light |
| 18:00 | 28 | Light |
| 19:00 | 27.5 | Light |
| 20:00 | 27 | Darkness |
| 21:00 | 26.5 | Darkness |
| 22:00 | 26 | Darkness |
| 23:00 | 25.5 | Darkness |
| 0:00 | 25 | Darkness |
| 1:00 | 24.5 | Darkness |
| 2:00 | 24 | Darkness |
| 3:00 | 23.5 | Darkness |
| 4:00 | 23 | Darkness |

8 days post hatching (i.e. when fish were transferred individually to glass jars), fish were randomly assigned to one of four experimental treatment groups (20 fish per group, Table 2.2): (1) temperature fluctuation without FLX ‘T(v)C’, (2) temperature fluctuation with 0.5 µg/L FLX ‘T(v)F’, (3) constant temperature without FLX ‘T(c)C’ and (4) constant temperature with 0.5 µg/L FLX ‘T(c)F’. This set-up was maintained until Day 59 (Table 2.2).

**Table 2.2.** Division of groups per condition (Day 9∼59)

|  | Temperature fluctuation | Control temperature |
| --- | --- | --- |
| Control water | T(v)C; 20 fish | T(c)C; 20 fish |
| FLX solution | T(v)F; 20 fish | T(c)F; 20 fish |

On Day 60, half of the fish of all experimental groups were exposed to 50 µg/L 3,4-DCA until the end of the experiment (Day 117), resulting in a total of 8 experimental groups with 10 replicate fish per group. The 8 groups were as follows (Table 2.3): (1) temperature fluctuation without FLX or 3,4-DCA ‘T(v)C’, (2) temperature fluctuation with 50 µg/L 3,4-DCA ‘T(v)P’, (3) temperature fluctuation with 0.5 µg/L FLX ‘T(v)F’, (4) temperature fluctuation with 0.5 µg/L FLX and 50 µg/L 3,4-DCA ‘T(v)F+P’, (5) constant temperature without FLX ‘T(c)C’, (6) constant temperature with 50 µg/L 3,4-DCA ‘T(c)P’, (7) constant temperature with 0.5 µg/L FLX ‘T(c)F’ and (8) constant temperature with 0.5 µg/L FLX and 50 µg/L 3,4-DCA ‘T(c)F+P’. This setting lasted until the end of the experiment (Day 117)

**Table 2.3.**
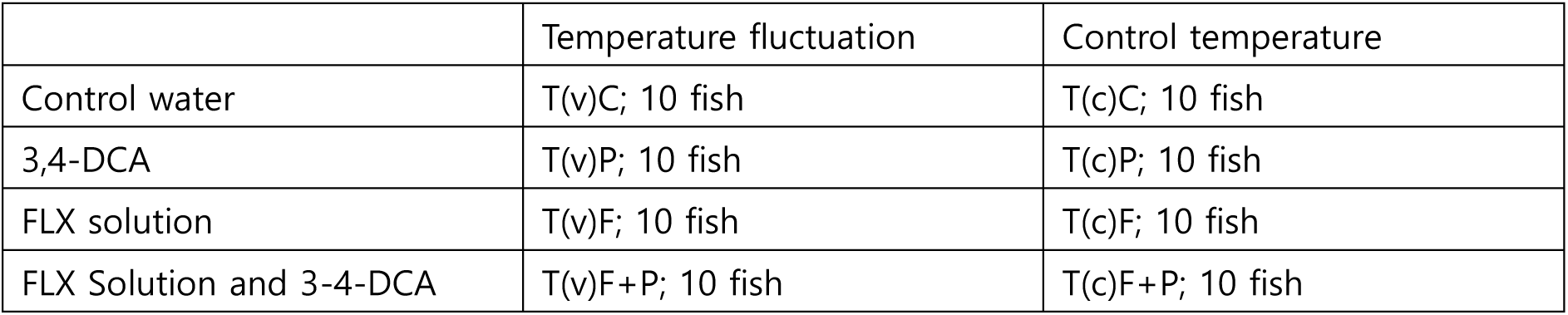
Division of groups per condition with initial sample size (Day 60∼117)

|  | Temperature fluctuation | Control temperature |
| --- | --- | --- |
| Control water | T(v)C; 10 fish | T(c)C; 10 fish |
| 3,4-DCA | T(v)P; 10 fish | T(c)P; 10 fish |
| FLX solution | T(v)F; 10 fish | T(c)F; 10 fish |
| FLX Solution and 3-4-DCA | T(v)F+P; 10 fish | T(c)F+P; 10 fish |

FLX and of 3,4-DCA concentrations were 0.5µg/L and 50 µg/L respectively. Within all treatment groups several fish died throughout the experiment and as a result this caused a slight reduction of sample size in each group by the end of experiment (Table 2.4).

**Table 2.4.**
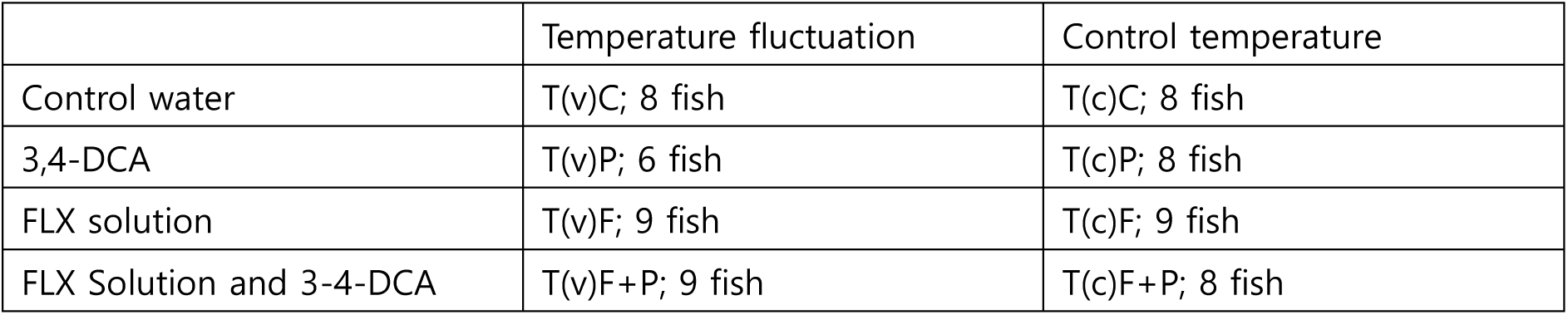
Division of groups per condition with final sample size (Day 117)

### 2.3. Preparation of solutions

FLX and 3,4-DCA treatments were applied twice a week to maintain a constant exposure dose throughout the experiment. This was done when water was refreshed during cleaning. The FLX stock solution (5 mg/L) had been prepared before the start of the experiment. This was added to the reconstituted water (600 µS/cm) and kept frozen. The frozen FLX stock solution was defrosted and added to the jars in the FLX groups to attain a 0.5 µg/L concentration. The 3,4-DCA stock solution was newly prepared one day before each cleaning session because of its typical fast degradation. The stock solution was stirred for 24 hours to ensure that the compound in the reconstituted water (600 µS/cm) was completely dissolved. The 3,4-DCA stock solution was added to the jars in the 3,4-DCA groups to reach a 50 µg/L concentration. Throughout the experiment water samples were taken from all treatments and frozen. These will serve to verify the actual realised FLX and 3,4-DCA concentrations via Mass Spectrometry. However, due to interruption of all experimental work in the light of Covid-19 measures the analyses could not yet be performed.

### 2.4. Endpoints

To examine the impacts of all stressors and their combinations, life-history and behavioural traits of all fish were monitored throughout the experiment. Life-history traits include body size, fecundity and behavioural traits include activity and boldness (analysed by open field test) as well as feeding behaviour (feeding test). C920 HD Pro Webcam (Logitech, 2020) and EthoVision XT Ver 9.0 (Noldus, 2020) were used to record and process, respectively open field test and feeding test data. Except for body size, all other traits were recorded only after fish reached maturity.

#### 2.4.1. Body size

The body size of Individual fish was measured once every two weeks starting on Day 22 (9 December 2019) until the end of the experiment (Figure 2.4). for a total of 7 repeated measures per individual.

**Figure 2.4.**
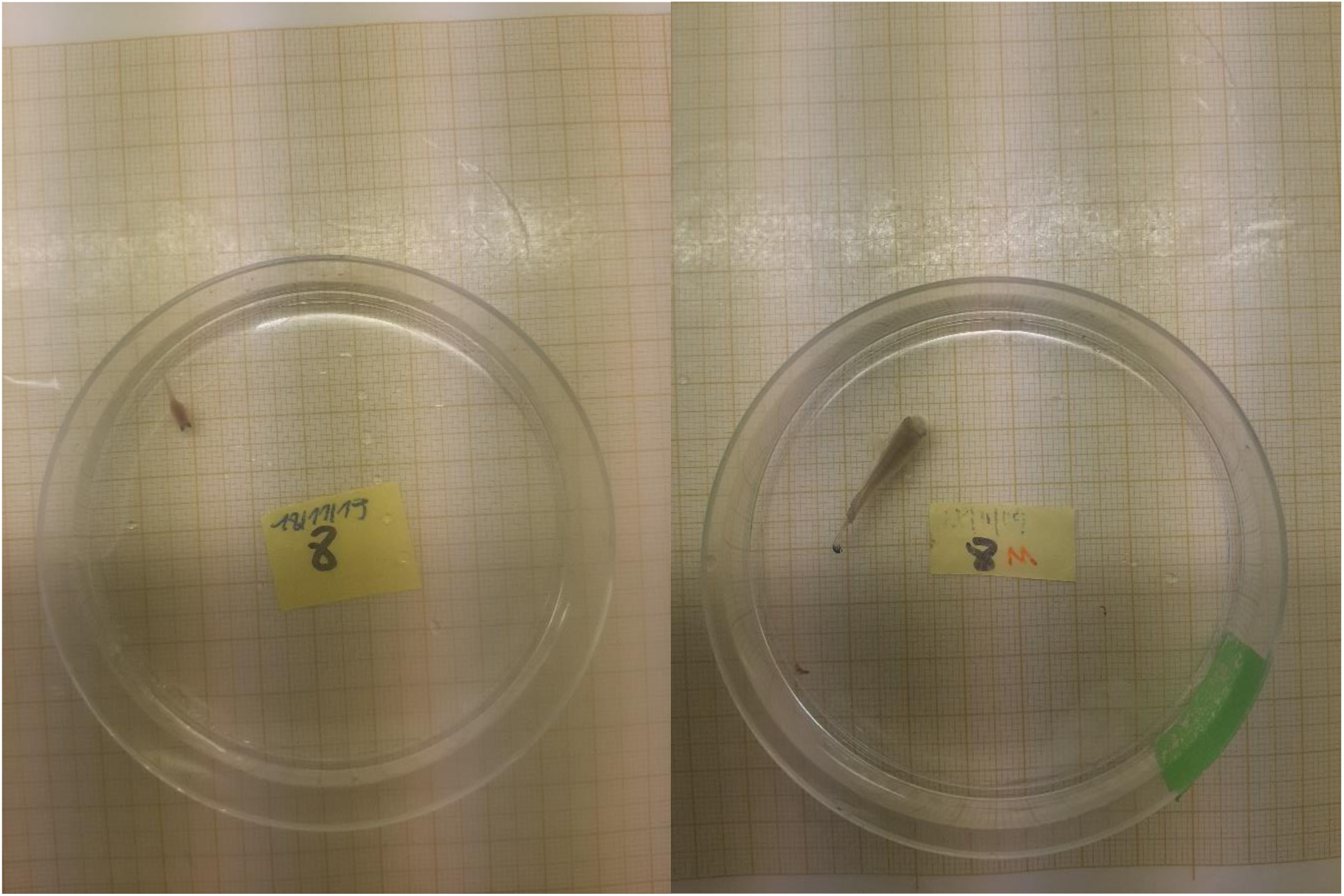
Body size measurement on Day 22 (first measurement; left) and on Day 113 (last measurement; right)

For body size measurements, each fish was transferred to a petri dish with a small amount of water. Photographs were standardised by taking a top view picture of the petri dish from a fixed distance. Size-calibrated, top-view (dorsal) photographs were taken and digitally analysed using ImageJ (Wayne Rasband, 2020). Body length was defined as the length from the tip of the snout to the end of the caudal fin.

#### 2.4.2. Open field test

An open field test was used to analyse horizontal locomotor activity and fish boldness. During the open field test, each fish was transferred individually to a test arena (17.4 cm x 11.2 cm x 11.5 cm). The test arena was filled with reconstituted water up to 2 cm (0.5L) to limit vertical movement. Fish were visually separated to prevent social interactions during the test and limit disturbance (Figure 2.5).

**Figure 2.5.**
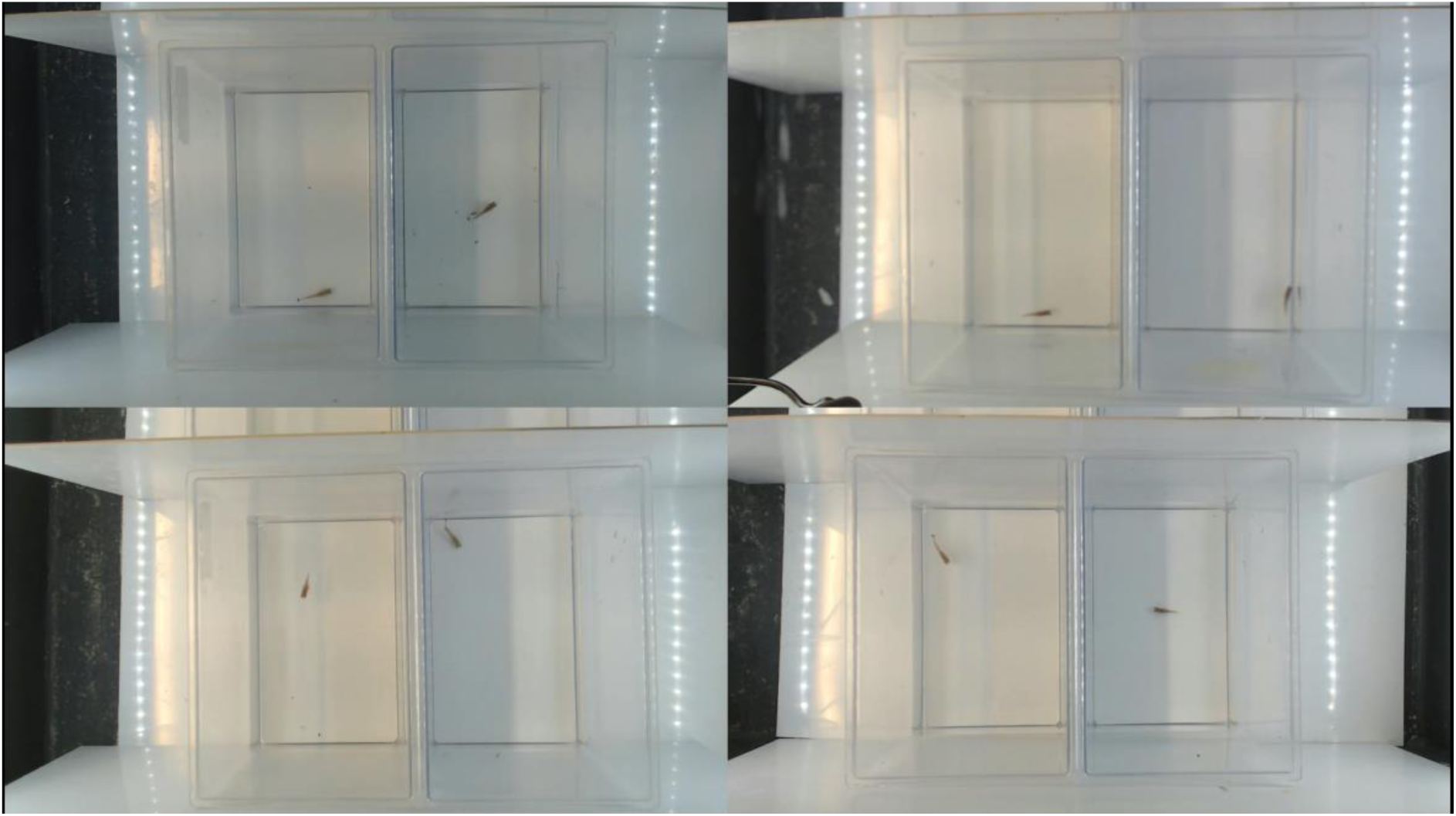
Open field test set-up

Each fish was allowed to acclimate for five minutes, after which fish activity was continuously recorded for 15 minutes. Open field tests were conducted 4 times (Day 50, Day 71, Day 85 and Day 106).

Locomotor activity was analysed by scoring 4 parameters: total distance travelled (cm), total time spent moving (s), maximum acceleration (cm/s²) and mean velocity (cm/s). In addition, each test arena was virtually divided in a centrum zone (50% middle of length and width) and a peripheral zone. Boldness was analysed by 3 parameters: latency time to enter the centre (s), cumulative distance between fish and centre of arena per frame (cm) and total distance moved in the centre (cm).

#### 2.4.3. Feeding test

The same test arenas and set-up as for the open field test were used for the feeding test. A fish was transferred to a test arena and allowed to acclimate for 5 mins, upon which fish were administered a standardized amount of frozen *Chironomus* larvae (Figure 2.6). Fish were filmed for 15 minutes, and the latency time to initiate feeding was taken as a measure for feeding propensity. Fish that failed to initiate feeding within this time were given the maximum value of 900sec (15 min). The recorded videos were analysed manually to identify the initial time to take the food. Feeding tests were conducted 4 times (Day 51, Day 72, Day 86 and Day 107).

**Figure 2.6.**
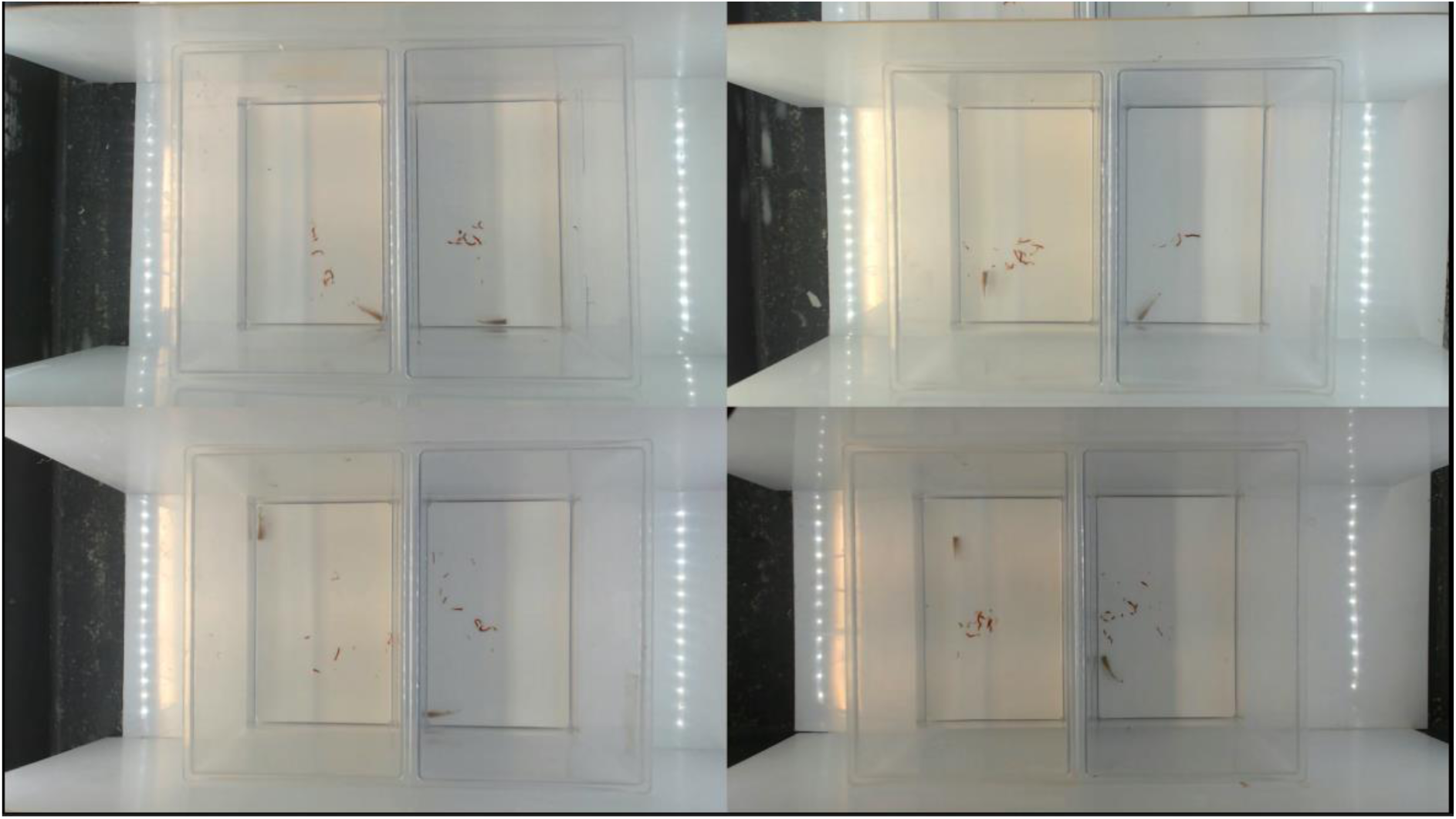
A trial of feeding test

#### 2.4.4. Fecundity test

The fecundity test was conducted twice a week from Day 74 until Day 117 for a total of 14 times per female fish. At the onset of each fecundity trial, one randomly selected male fish and one randomly selected female fish was paired in a 1L tank (Figure 2.7). The tank was filled with 2 cm of fine (<500µm) sand substrate (to allow for eggs deposition) and 1L of the reconstituted water. All pairs were allowed to spawn for 3 hours per trial.

**Figure 2.7.**
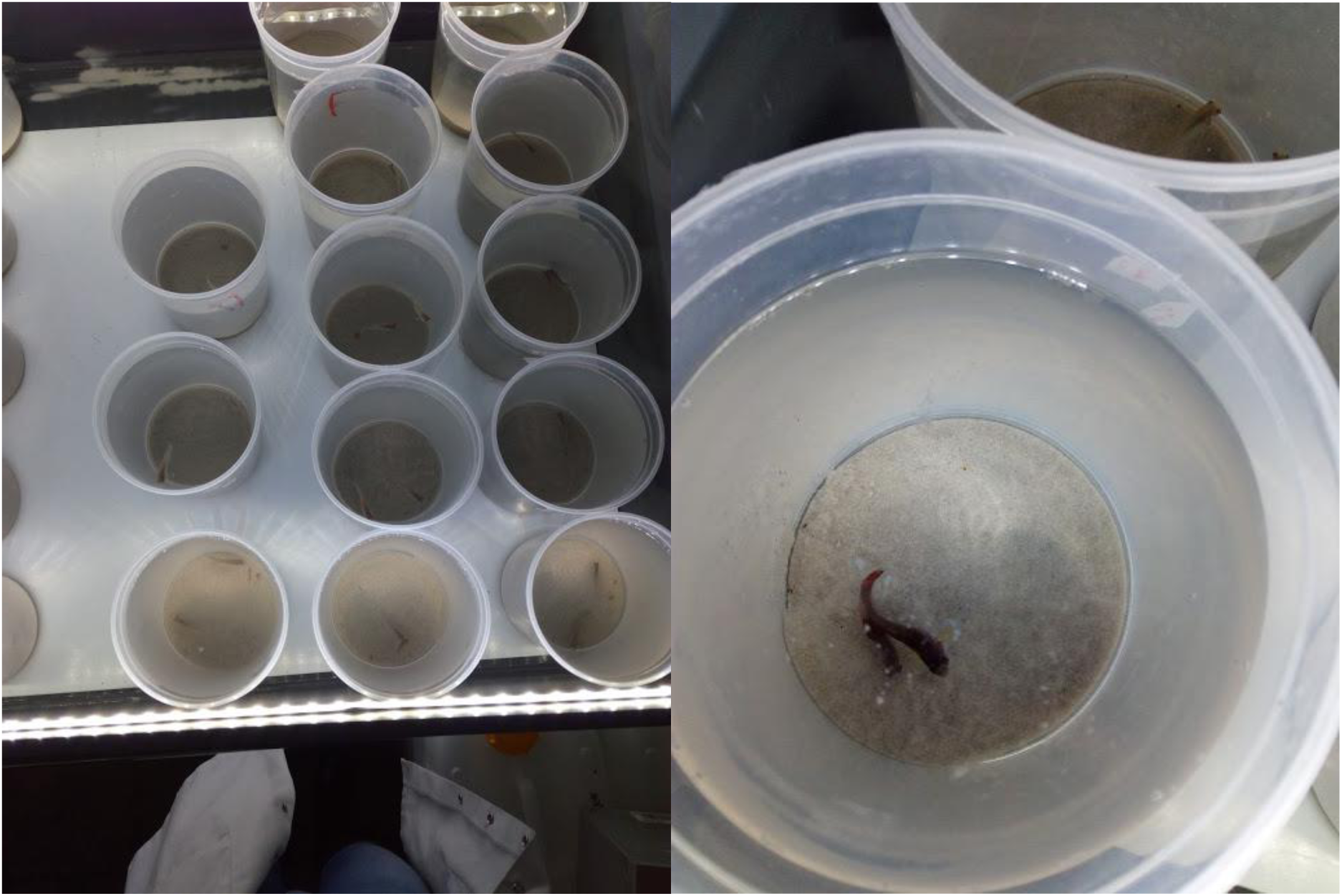
Fish were paired for 3h in a 1L jar with fine sand to spawn (Van Hooreweghe, 2019)

Afterwards, each individual fish was transferred back to its own jar. A 1mm grid sieve was used to retrieve eggs from the sand. The number of eggs was counted, and the eggs were subsequently stored on top of moist peat (Jiffy-7c) in a polystyrene petri dish and transferred to a dark incubator at 17°C for long term storage.

### 2.5. Data analysis

Statistical analyses were performed in R i386 4.0.0 (R Development Core Team, 2020). Significant level was 0.05 throughout all tests conducted and model assumptions of each variable (condition) were graphically verified. Linear mixed model (LMM) (lme4 package; Bates et al., 2017) were used to analyse all behavioural endpoints (activity, boldness and feeding traits). Gaussian error distribution was assumed for all endpoints including behaviour endpoints, body size and fecundity. A factor defined ‘condition’ (Table 2.4) consists of combinations from temperature, FLX, 3,4-DCA and the combination of FLX and 3,4-DCA. Acceleration, distance moved, time moved, trial (each recording), conditions and sex (female and male) were modelled as fixed factors while ID was modelled as a random factor for assessing activity and boldness. Time taken, trial, conditions and sex were modelled as fixed factors while ID was modelled as a random factor for assessing feeding responses. Post-hoc tests were subsequently performed (lsmeans package) to verify the effects of combinations of 3,4-DCA and temperature on the response variables (Lenth, 2015). ANOVA tests were done to analyse body size, measured at 8 times, 2-week-intervals (each measurement was counted as a ‘trial’), and fecundity, number of collected eggs (each test was counted as a ‘trial’). Trial, conditions and sex were fixed factors and ID was a random factor in the model evaluating body size effects of the tested conditions. Number of eggs per female, trial, conditions and sex were fixed factors and ID was a random factor in fecundity. Type 3 Wald chi-square tests from a car package were used to test significance of fixed factors (John et al., 2020).

## 3. Results

### 3.1. Effects on feeding behaviour

The latency time to initiate feeding (Figure 3.1) was shorter in the FLX group (125.511 ± 27.770 s) than in the control group (222.600 ± 23.682 s) during the feeding test. 3,4-DCA and temperature fluctuation had no influence on the latency time to initiate feeding (Table 3.1).

**Figure 3.1.**
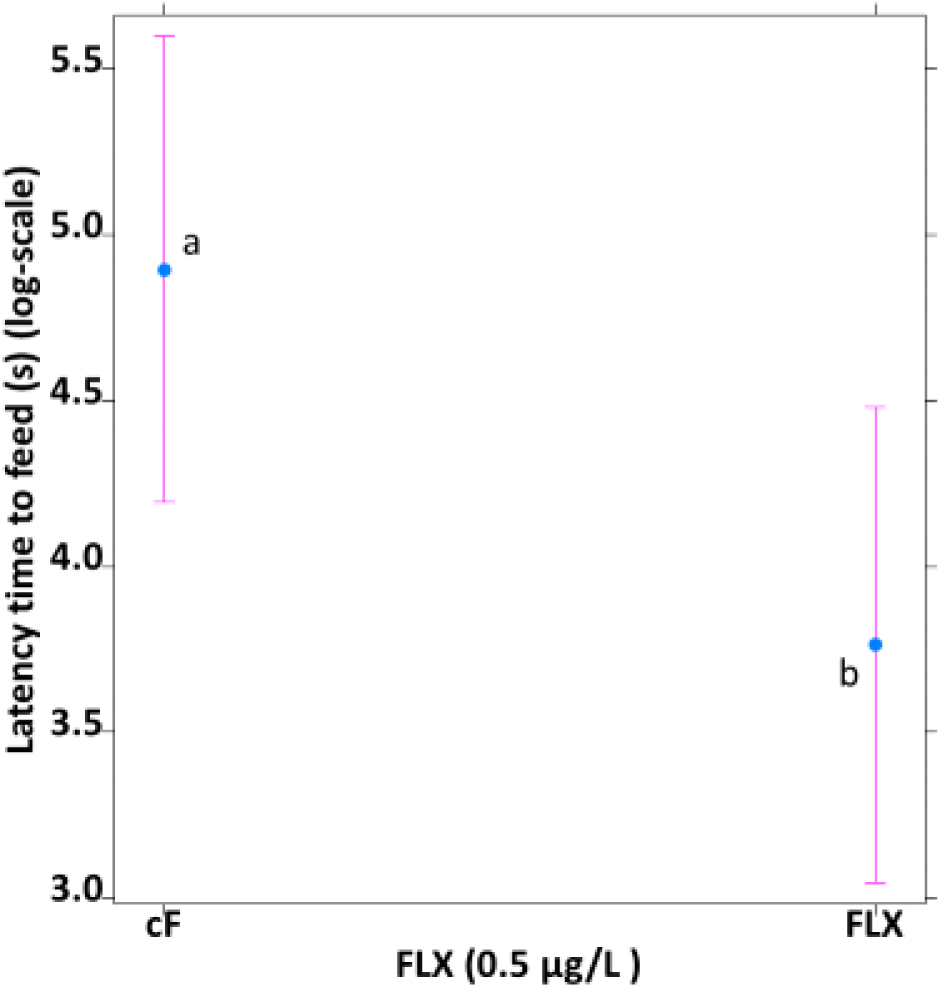
Impact of FLX-exposure on *N. furzeri* feeding behaviour. Average latency time (in seconds, logarithmic scale) to start feeding in the feeding test for control fish (cF) and fish exposed to 0.5 µg/L FLX (FLX). Whiskers delineate the upper and lower 95% confidence limit. Letters indicate significant differences based on Tukey-corrected post-hoc tests.

**Table 3.1.** The results of the LMM to analyse the latency time to initiate feeding in relation to the tested temperature, FLX and 3,4-DCA conditions.

| <u>Condition</u> | <u>$\chi^2$</u> | <u>p-value</u> |
| --- | --- | --- |
| FLX | 18.9326 | < **0.001** |
| 3,4-DCA | 3.0613 | 0.08018 |
| Temperature | 0.0042 | 0.94833 |
| Sex | 3.0287 | 0.08181 |
| Body Size | 0.7997 | 0.37118 |
| FLX:3,4-DCA | 3.5449 | 0.05973 |
| FLX:Temperature | 0.1349 | 0.71337 |
| 3,4-DCA:Temperature | 4.9140 | **0.02664** |
| FLX:Sex | 0.5096 | 0.47533 |
| 3,4-DCA:Sex | 0.8088 | 0.36849 |
| Temperature:Sex | 0.0395 | 0.84253 |
| FLX:3,4-DCA:Temperature | 0.0009 | 0.97666 |
| FLX:3,4-DCA:Sex | 0.6804 | 0.40945 |
| FLX:Temperature:Sex | 1.8487 | 0.17393 |
| 3,4-DCA:Temperature:Sex | 0.4677 | 0.49405 |
| FLX:3,4-DCA:Temperature:Sex | 0.3703 | 0.54285 |

The combination of 3,4-DCA and temperature affected the latency time to initiate feeding as well (Table 3.2). The temperature fluctuation group was not affected by this combination. However, in the control temperature group, the 3,4-DCA group had a longer latency time to start feeding than the control group (Figure 3.2).

**Table 3.2.**
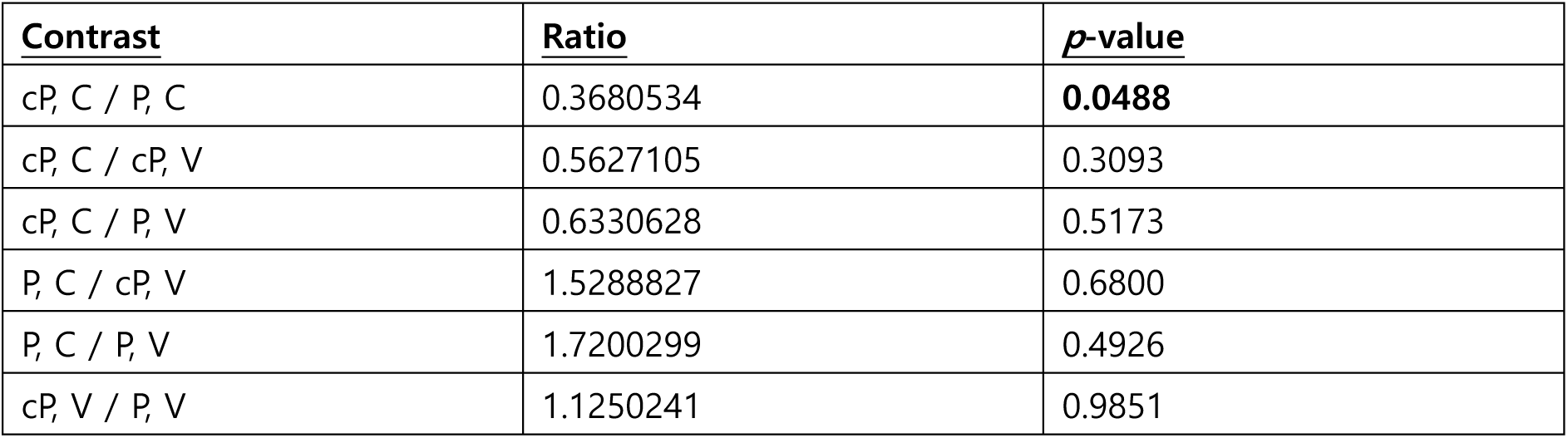
The result of the Post-hoc on the combination of 3,4-DCA and temperature to analyse the latency time to start feeding.

| **Contrast** | **Ratio** | **p-value** |
| --- | --- | --- |
| cP, C / P, C | 0.3680534 | **0.0488** |
| cP, C / cP, V | 0.5627105 | 0.3093 |
| cP, C / P, V | 0.6330628 | 0.5173 |
| P, C / cP, V | 1.5288827 | 0.6800 |
| P, C / P, V | 1.7200299 | 0.4926 |
| cP, V / P, V | 1.1250241 | 0.9851 |

**Figure 3.2.**
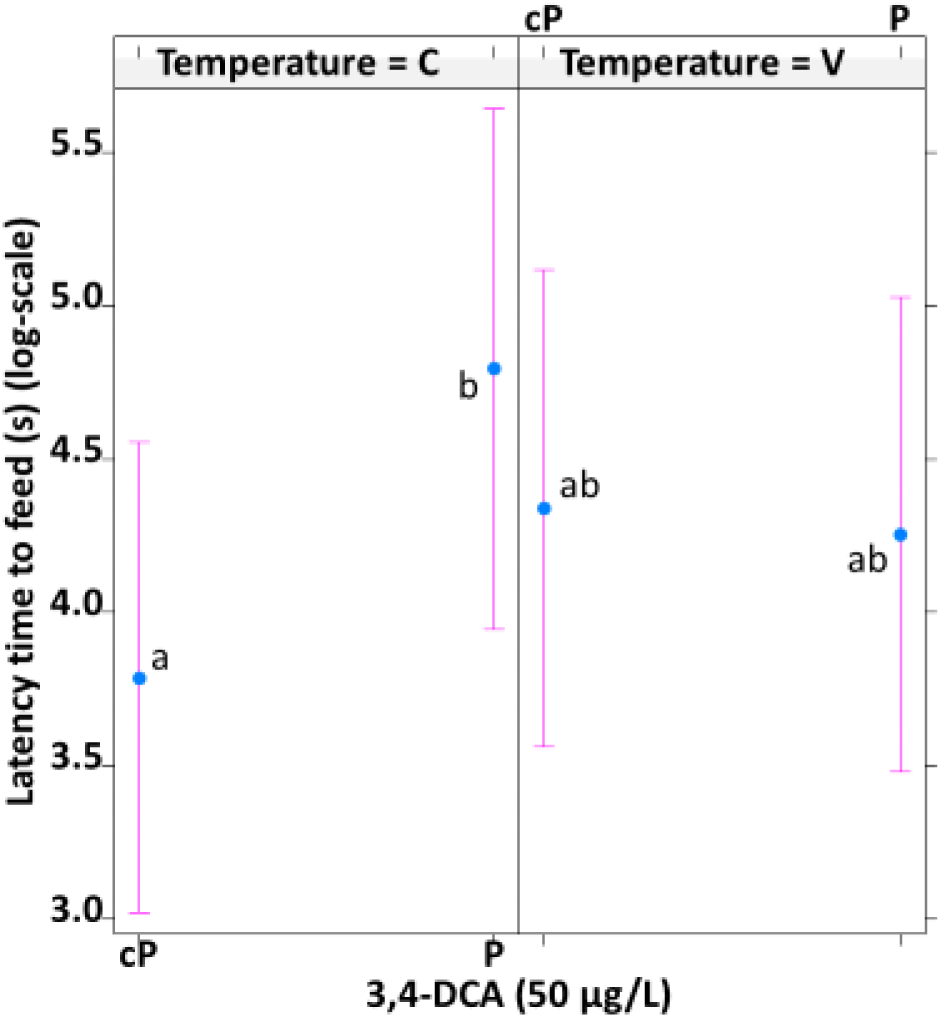
Impact of pesticide-exposure (3,4-DCA) on *N. furzeri* feeding behaviour under different temperature scenarios. Average latency time (in seconds, logarithmic scale) to start feeding in the feeding test for control fish (cP) and fish exposed to 50 µg/L 3,4-DCA (P) under constant (C) and fluctuating (F) temperature conditions. Whiskers delineate the upper and lower 95% confidence limit. Letters indicate significant differences based on Tukey-corrected post-hoc tests.

### 3.2. Effects on activity

#### 3.2.1. Total distance travelled

Fish of the 3,4-DCA group (32729.403 ±SD: 0.05) travelled a larger distance compared to the control group (32127.081 ± 253.012 cm) during the open field test (Figure 3.3). The total distance travelled was not influenced by FLX and temperature. Body size showed observable relevance to the total distance travelled but that effect was not significant (Table 3.3).

**Figure 3.3.**
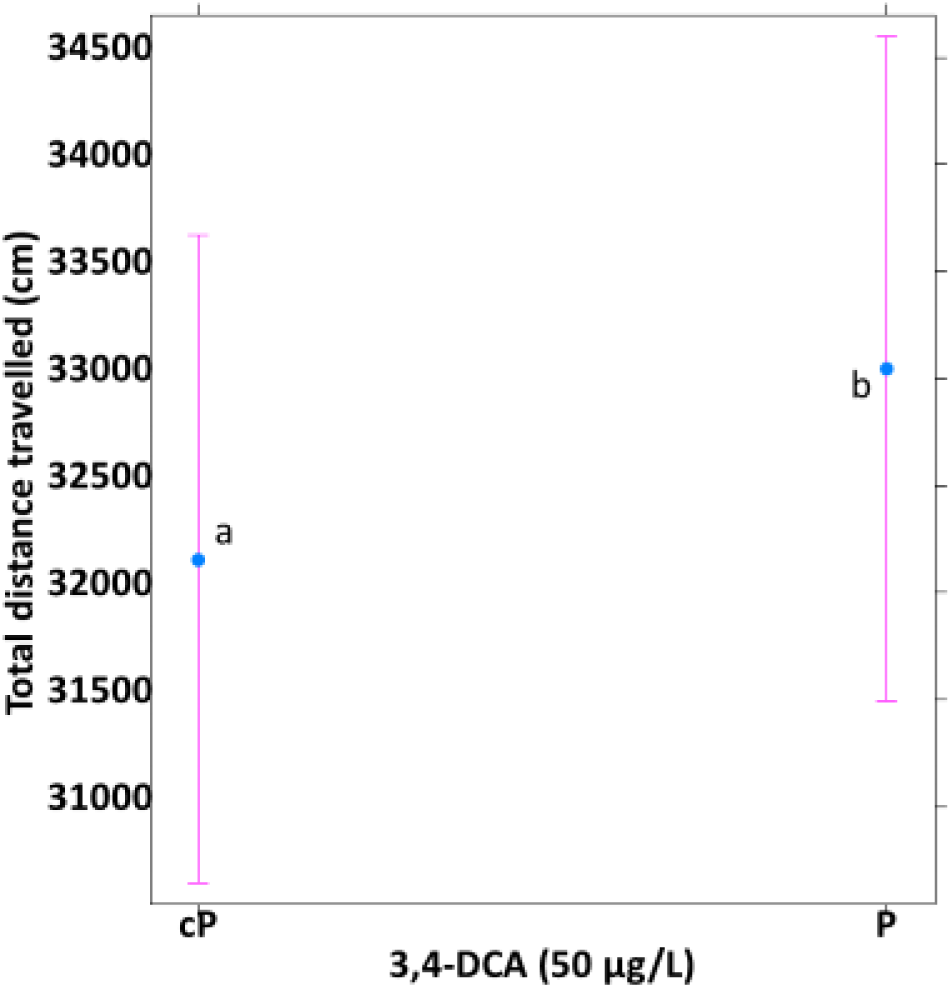
Impact of pesticide-exposure (3,4-DCA) on *N. furzeri* activity. Total distance travelled (in centimetres) in the open field test for control fish (cP) and fish exposed to 50 µg/L 3,4-DCA (P). Whiskers delineate the upper and lower 95% confidence limit. Letters indicate significant differences based on Tukey-corrected post-hoc tests.

**Table 3.3.** The result of the LMM to analyse the total distance travelled in relation to the tested temperature, FLX and 3,4-DCA conditions.

| Condition | $\chi^2$ | *p*-value |
| --- | --- | --- |
| FLX | 1.7035 | 0.19183 |
| 3,4-DCA | 4.9760 | **0.02570** |
| Temperature | 0.3362 | 0.56206 |
| Sex | 0.7614 | 0.38288 |
| Body Size | 3.6029 | 0.05768 |
| FLX:3,4-DCA | 0.3852 | 0.53484 |
| FLX:Temperature | 0.2842 | 0.59393 |
| 3,4-DCA:Temperature | 0.0125 | 0.91106 |
| FLX:Sex | 0.3599 | 0.54858 |
| 3,4-DCA:Sex | 2.5209 | 0.11235 |
| Temperature:Sex | 0.9310 | 0.33460 |
| FLX:3,4-DCA:Temperature | 1.0166 | 0.31333 |
| FLX:3,4-DCA:Sex | 0.0288 | 0.86529 |
| FLX:Temperature:Sex | 0.0404 | 0.84064 |
| 3,4-DCA:Temperature:Sex | 0.0806 | 0.77655 |
| FLX:3,4-DCA:Temperature:Sex | 1.8256 | 0.17665 |

#### 3.2.2. Total time spent moving

Fish from the FLX group (372.768 ± 14.047 s) moved significantly longer compared to control fish (275.150 ± 14.648 s) during the open field test (Figure 3.4). 3,4-DCA and temperature did not affect the total time spent moving (Table 3.4).

**Figure 3.4.**
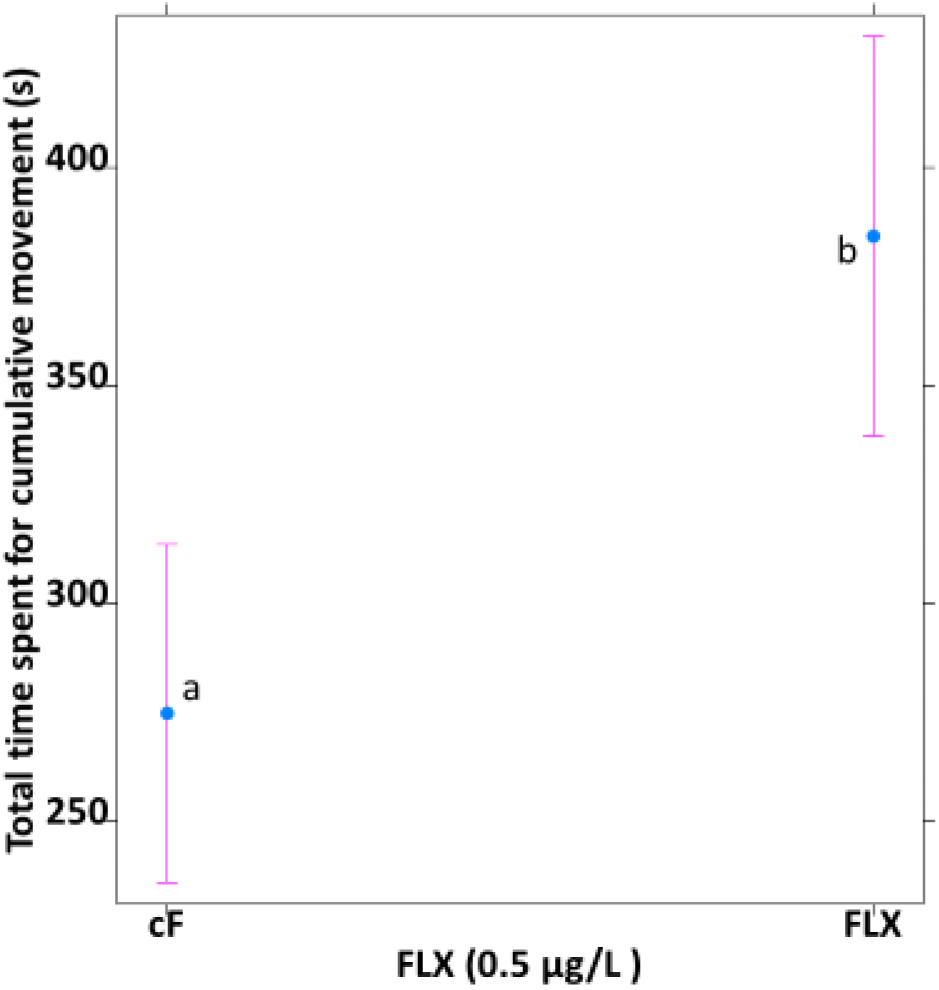
Impact of FLX-exposure on *N. furzeri* activity. Total time spent for cumulative movement (in seconds) in the open field test for control fish (cF) and fish exposed to 0.5 µg/L FLX (FLX). Whiskers delineate the upper and lower 95% confidence limit. Letters indicate significant differences based on Tukey-corrected post-hoc tests.

**Table 3.4.** The result of the LMM to analyse the total time spent for cumulative movement in relation to the tested temperature, FLX and 3,4-DCA conditions.

| <u>Condition</u> | <u>$\chi^2$</u> | <u>*p*-value</u> |
| --- | --- | --- |
| FLX | 12.3350 | < **0.001** |
| 3,4-DCA | 0.2774 | 0.5984224 |
| Temperature | 2.5342 | 0.1114026 |
| Sex | 2.0039 | 0.1568955 |
| Body Size | 0.4604 | 0.4974278 |
| FLX:3,4-DCA | 1.3788 | 0.2403100 |
| FLX:Temperature | 2.6786 | 0.1017067 |
| 3,4-DCA:Temperature | 0.1694 | 0.6806259 |
| FLX:Sex | 1.0081 | 0.3153655 |
| 3,4-DCA:Sex | 0.0371 | 0.8471761 |
| Temperature:Sex | 2.8249 | 0.0928110 |
| FLX:3,4-DCA:Temperature | 0.4450 | 0.5047339 |
| FLX:3,4-DCA:Sex | 0.8404 | 0.3592902 |
| FLX:Temperature:Sex | 0.0628 | 0.8021486 |
| 3,4-DCA:Temperature:Sex | 0.0747 | 0.7845616 |
| FLX:3,4-DCA:Temperature:Sex | 0.0431 | 0.8355728 |

#### 3.2.3. Maximum acceleration

There was no specific condition that influenced the maximum acceleration of individuals during the open field test (Table 3.5). There was a slight difference of the maximum acceleration depending on sex, with males being faster than female, but this was not significant (Figure 3.5).

**Table 3.5.** The result of the LMM to analyse the maximum acceleration in relation to the tested temperature, FLX and 3,4-DCA conditions.

| <u>Condition</u> | <u>$\chi^2$</u> | <u>p-value</u> |
| --- | --- | --- |
| FLX | 2.3434 | 0.1258117 |
| 3,4-DCA | 0.0153 | 0.9013987 |
| Temperature | 0.4625 | 0.4964493 |
| Sex | 0.5612 | 0.4537907 |
| Body Size | 1.0954 | 0.2952782 |
| FLX:3,4-DCA | 0.9176 | 0.3381000 |
| FLX:Temperature | 0.0248 | 0.8748535 |
| 3,4-DCA:Temperature | 0.0602 | 0.8061687 |
| FLX:Sex | 2.0673 | 0.1504866 |
| 3,4-DCA:Sex | 1.3164 | 0.2512464 |
| Temperature:Sex | 1.0183 | 0.3129187 |
| FLX:3,4-DCA:Temperature | 1.5148 | 0.2184138 |
| FLX:3,4-DCA:Sex | 0.2179 | 0.6406559 |
| FLX:Temperature:Sex | 0.0331 | 0.8556003 |
| 3,4-DCA:Temperature:Sex | 0.0000 | 0.9995004 |
| FLX:3,4-DCA:Temperature:Sex | 0.3243 | 0.5690195 |

**Figure 3.5.**
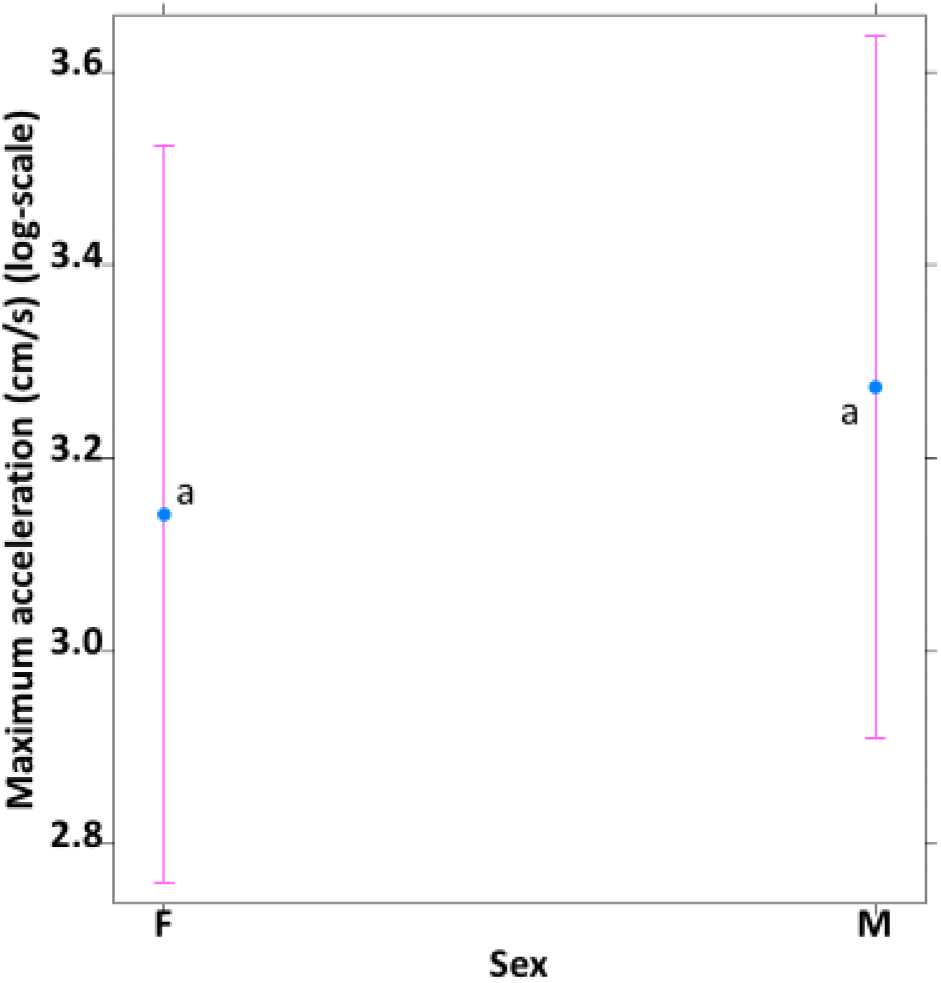
Impact of sex on *N. furzeri* activity. Maximum acceleration (cm/s, logarithmic scale) in the open field test for female fish (F) and male fish (M). Whiskers delineate the upper and lower 95% confidence limit. Letters indicate significant differences based on Tukey-corrected post-hoc tests.

#### 3.2.4. Mean velocity

There was a strong effect of FLX on the mean velocity of individuals as fish from the FLX group (1.810 ± 0.064 cm/s) were notably faster than individuals from the control group (1.345 ± 0.060 cm/s) during the open field test (Figure 3.6). The mean velocity was not impacted by 3,4-DCA or temperature regimes (Table 3.6).

**Figure 3.6.**
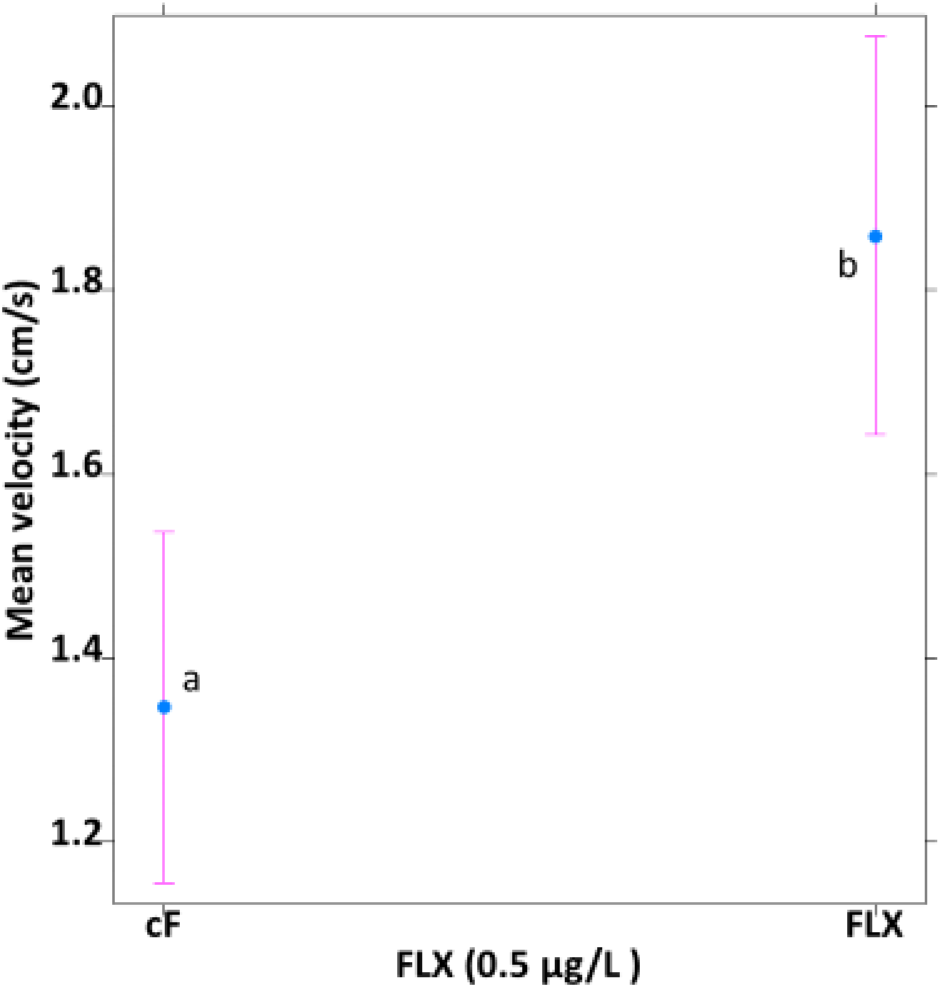
Impact of FLX-exposure on *N. furzeri* activity. Mean velocity (in cm/s) in the open field test for control fish (cF) and fish exposed to 0.5 µg/L FLX (FLX) under constant (C) and fluctuating (F) temperature conditions. Whiskers delineate the upper and lower 95% confidence limit. Letters indicate significant differences based on Tukey-corrected post-hoc tests.

**Table 3.6.** The result of the LMM to analyse the mean velocity in relation to the tested temperature, FLX and 3,4-DCA conditions.

| <u>Condition</u> | <u>$\chi^2$</u> | <u>*p*-value</u> |
| --- | --- | --- |
| FLX | 15.4515 | < **0.001** |
| 3,4-DCA | 0.3720 | 0.54189 |
| Temperature | 2.1151 | 0.14586 |
| Sex | 2.2290 | 0.13544 |
| Body Size | 0.6267 | 0.42855 |
| FLX:3,4-DCA | 1.2565 | 0.26231 |
| FLX:Temperature | 3.5094 | 0.06102 |
| 3,4-DCA:Temperature | 0.2450 | 0.62064 |
| FLX:Sex | 0.6158 | 0.43262 |
| 3,4-DCA:Sex | 0.0977 | 0.75466 |
| Temperature:Sex | 2.5480 | 0.11044 |
| FLX:3,4-DCA:Temperature | 0.3799 | 0.53768 |
| FLX:3,4-DCA:Sex | 0.8860 | 0.34658 |
| FLX:Temperature:Sex | 0.0907 | 0.76333 |
| 3,4-DCA:Temperature:Sex | 0.1887 | 0.66402 |
| FLX:3,4-DCA:Temperature:Sex | 0.0242 | 0.87634 |

### 3.3. Effects on boldness

#### 3.3.1. Latency time to enter the centre zone

The latency time to enter the centre zone was mainly dependent on the temperature treatment. During the open field test, fish from the temperature fluctuation group (146.451 ± 4.764 s) spent longer to enter the zone than fish from the control temperature group (62.177 ± 4.287 s) (Figure 3.7). FLX and 3,4-DCA did not influence the latency time to enter the centre zone (Table 3.7).

**Figure 3.7.**
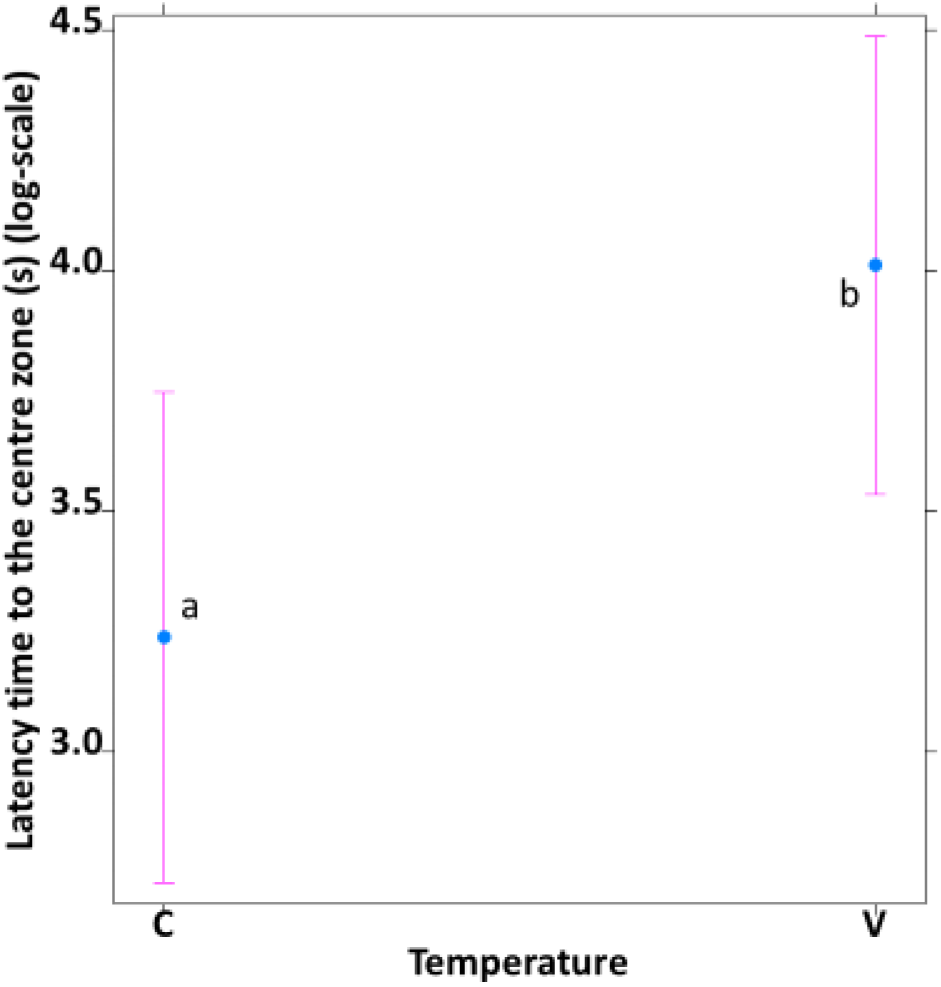
Impact of temperature variation exposure on *N. furzeri* boldness. Latency time to the centre zone (in seconds, logarithmic scale) in the open field test for control fish (C) and fish exposed to temperature variation (V). Whiskers delineate the upper and lower 95% confidence limit. Letters indicate significant differences based on Tukey-corrected post-hoc tests.

**Table 3.7.** The result of the LMM to analyse the latency time to the centre zone in relation to the tested temperature, FLX and 3,4-DCA conditions.

| <u>Condition</u> | <u>$\chi^2$</u> | <u>p-value</u> |
| --- | --- | --- |
| FLX | 0.0032 | 0.955118 |
| 3,4-DCA | 0.5317 | 0.465875 |
| Temperature | 10.8687 | < **0.001** |
| Sex | 0.2482 | 0.618324 |
| Body Size | 0.0057 | 0.939773 |
| FLX:3,4-DCA | 1.1835 | 0.276652 |
| FLX:Temperature | 1.8610 | 0.172515 |
| 3,4-DCA:Temperature | 0.6493 | 0.420357 |
| FLX:Sex | 0.0552 | 0.814277 |
| 3,4-DCA:Sex | 1.1641 | 0.280613 |
| Temperature:Sex | 0.0155 | 0.900890 |
| FLX:3,4-DCA:Temperature | 0.5359 | 0.464133 |
| FLX:3,4-DCA:Sex | 0.0081 | 0.928073 |
| FLX:Temperature:Sex | 0.1055 | 0.745329 |
| 3,4-DCA:Temperature:Sex | 0.0054 | 0.941537 |
| FLX:3,4-DCA:Temperature:Sex | 0.4701 | 0.492961 |

#### 3.3.2. Cumulative distance between fish and centre of arena per frame

The cumulative distance between fish and centre of arena per frame strongly influenced by FLX. The average of individuals from the FLX group (1589.861 ± 59.341 cm) moved more than the average of individuals from the control group (1230.616 ± 50.329 cm) during the open field test (Figure 3.8). 3,4-DCA and temperature did not impact the total distance moved in the centre zone (Table 3.8).

**Figure 3.8.**
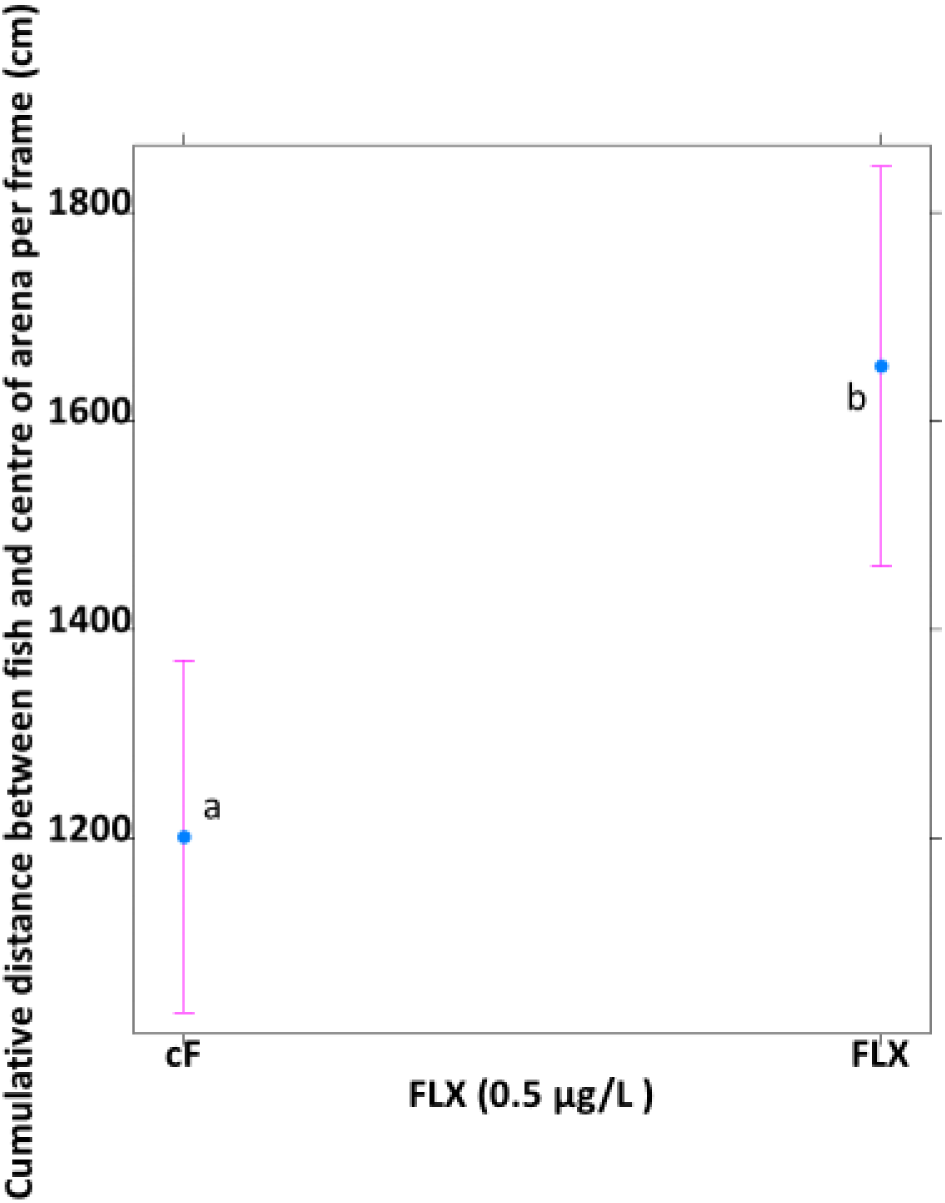
Impact of FLX-exposure on *N. furzeri* boldness. cumulative distance between fish and the centre of the arena per frame (in centimetre) in the open field test for control fish (cF) and fish exposed to 0.5 µg/L FLX (FLX). Whiskers delineate the upper and lower 95% confidence limit. Letters indicate significant differences based on Tukey-corrected post-hoc tests.

**Table 3.8.** The result of the LMM to analyse the total distance moved in the centre zone in relation to the tested temperature, FLX and 3,4-DCA conditions.

| **Condition** | **$\chi^2$** | ***p*-value** |
| --- | --- | --- |
| FLX | 15.2730 | < **0.001** |
| 3,4-DCA | 0.3055 | 0.58046 |
| Temperature | 2.0152 | 0.15573 |
| Sex | 2.1240 | 0.14500 |
| Body Size | 0.4290 | 0.51248 |
| FLX:3,4-DCA | 1.1854 | 0.27626 |
| FLX:Temperature | 3.5075 | 0.06109 |
| 3,4-DCA:Temperature | 0.2827 | 0.59494 |
| FLX:Sex | 0.5525 | 0.45732 |
| 3,4-DCA:Sex | 0.1145 | 0.73509 |
| Temperature:Sex | 2.6655 | 0.10255 |
| FLX:3,4-DCA:Temperature | 0.4213 | 0.51631 |
| FLX:3,4-DCA:Sex | 0.9219 | 0.33697 |
| FLX:Temperature:Sex | 0.0990 | 0.75305 |
| 3,4-DCA:Temperature:Sex | 0.1900 | 0.66295 |
| FLX:3,4-DCA:Temperature:Sex | 0.0278 | 0.86760 |

#### 3.3.3. Total time spent in the centre zone

The total time spent in the centre zone was especially influenced by temperature regime. Fish from the control temperature group (38.991 ± 2.413 s) stayed in the centre zone for a longer amount of time than the fish from the temperature fluctuation group (38.932 ± 2.022 s) during the open field test (Figure 3.9). FLX and 3,4-DCA did not significantly affect the total time spent in the centre zone (Table 3.9).

**Figure 3.9.**
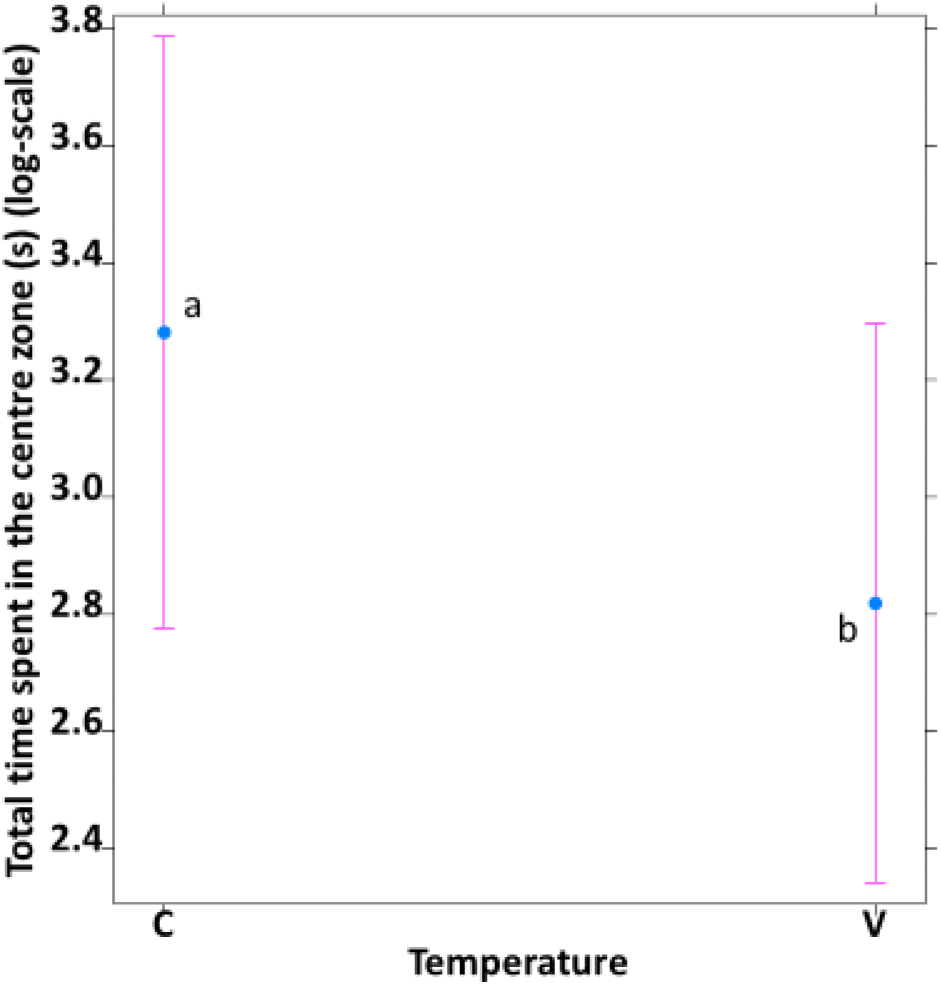
Impact of temperature variation on *N. furzeri* boldness. Total time spent in the centre zone (in seconds, logarithmic scale) in the open field test for control fish (C) and fish exposed to temperature variation (V). Whiskers delineate the upper and lower 95% confidence limit. Letters indicate significant differences based on Tukey-corrected post-hoc tests.

**Table 3.9.** The result of the LMM to analyse the total time spent in the centre zone in relation to the tested temperature, FLX and 3,4-DCA conditions.

| <u>Condition</u> | <u>$\chi^2$</u> | <u>*p*-value</u> |
| --- | --- | --- |
| FLX | 0.0032 | 0.95483 |
| 3,4-DCA | 0.9335 | 0.33395 |
| Temperature | 4.5829 | 0.03229 |
| Sex | 0.2427 | 0.62223 |
| Body Size | 0.0184 | 0.89209 |
| FLX:3,4-DCA | 3.2554 | 0.07119 |
| FLX:Temperature | 0.0781 | 0.77988 |
| 3,4-DCA:Temperature | 0.0179 | 0.89370 |
| FLX:Sex | 1.1249 | 0.28886 |
| 3,4-DCA:Sex | 0.0049 | 0.94412 |
| Temperature:Sex | 0.6960 | 0.40412 |
| FLX:3,4-DCA:Temperature | 0.2034 | 0.65202 |
| FLX:3,4-DCA:Sex | 0.5278 | 0.46753 |
| FLX:Temperature:Sex | 0.1326 | 0.71578 |
| 3,4-DCA:Temperature:Sex | 0.3471 | 0.55577 |
| FLX:3,4-DCA:Temperature:Sex | 1.0508 | 0.30532 |

### 3.4. Body size

Body size was very strongly sex dependent with males (26.534 ± 1.244 cm) being larger than females (21.909 ± 1.421 cm) (Figure 3.10). FLX, 3,4-DCA and temperature did not influence the body size (Table 3.10).

**Figure 3.10.**
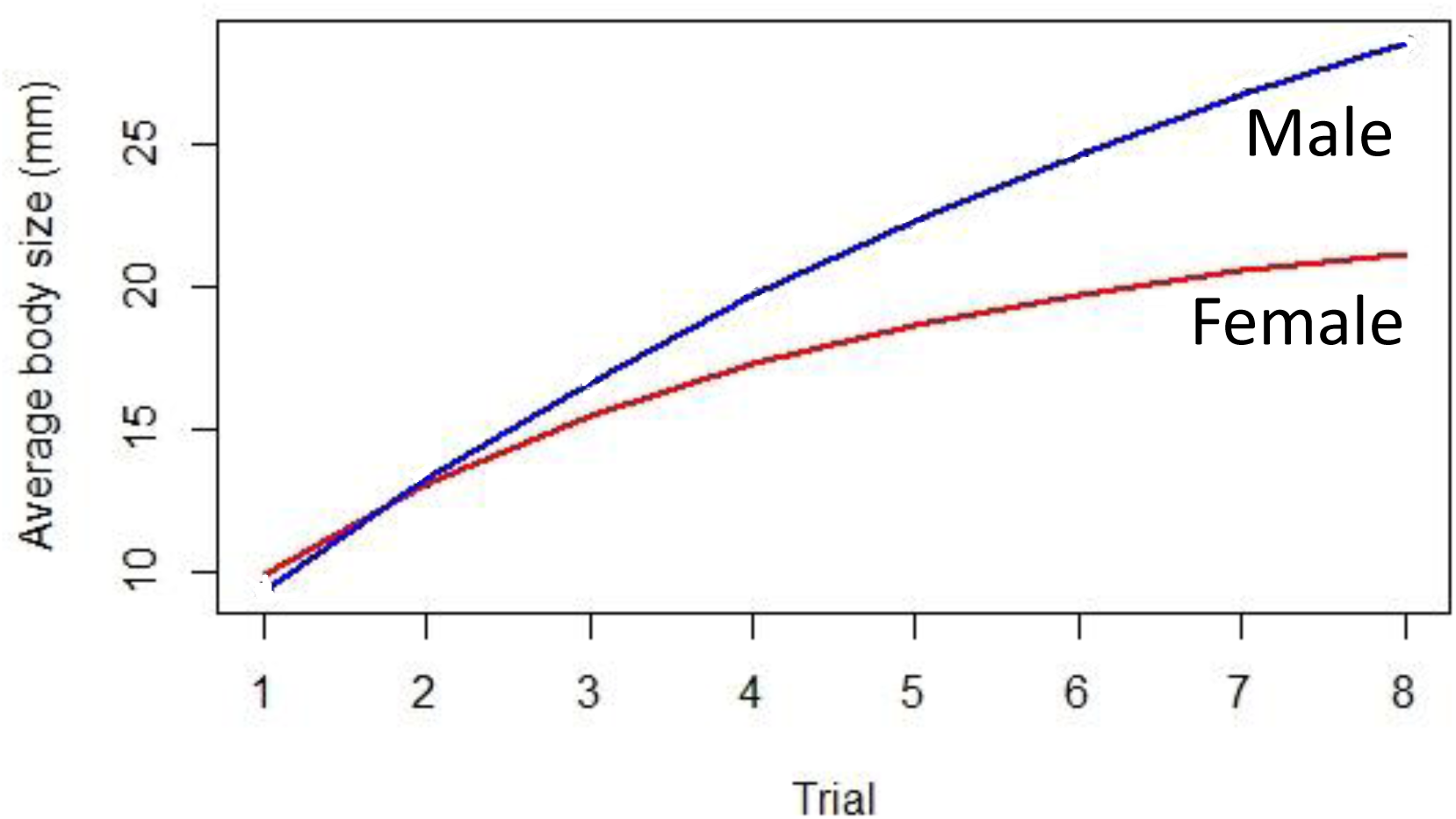
Impact of sex on *N. furzeri* body size. Average body size (in millimetres) in the body size measurement for female fish (red line) and male fish (blue line). Lines delineate the upper and lower 95% confidence limit.

**Figure 3.11.**
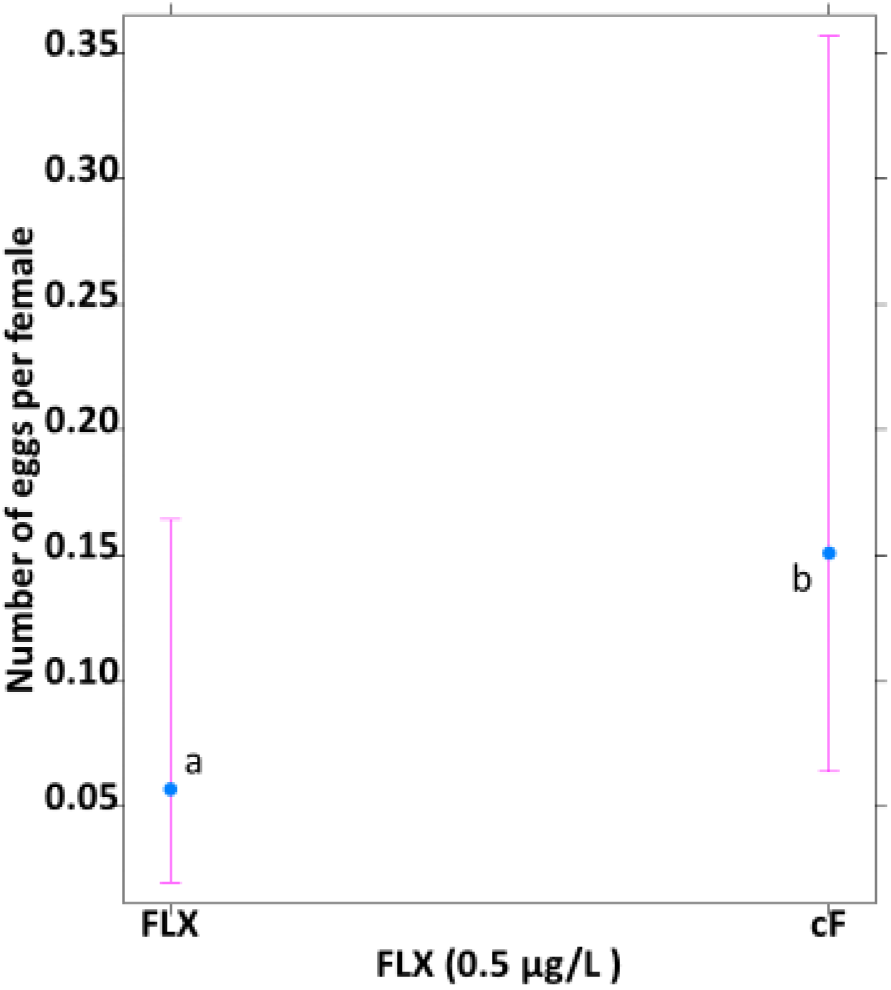
Impact of FLX-exposure on *N. furzeri* fecundity. Number of eggs per female in the fecundity test for control fish (cF) and fish exposed to 0.5 µg/L FLX (FLX). Whiskers delineate the upper and lower 95% confidence limit. Letters indicate significant differences based on Tukey-corrected post-hoc tests.

**Figure 3.12.**
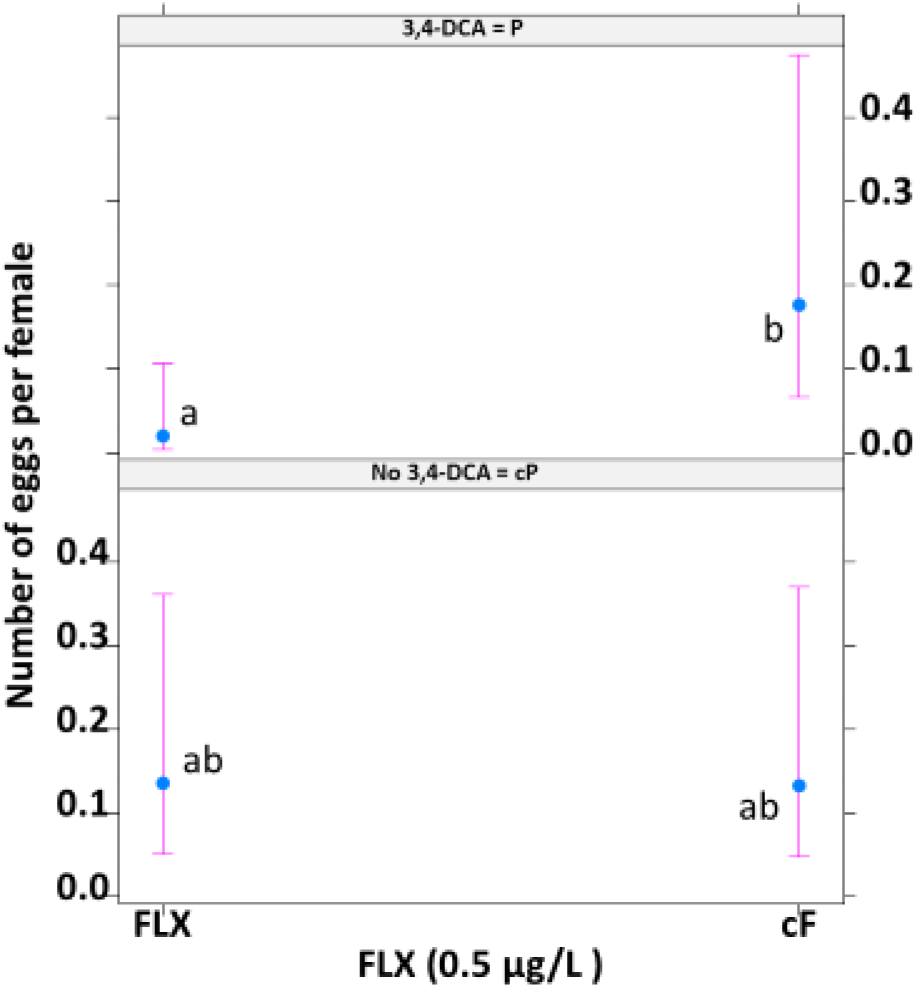
Impact of FLX-exposure on *N. furzeri* fecundity under different pesticide-exposure (3,4-DCA) scenarios. Number of eggs per female in the fecundity test for control fish (cF) and fish exposed to 0.5 µg/L FLX (FLX) under no 3,4-DCA-exposure (cP) and 3,4-DCA-exposure (P) conditions. Whiskers delineate the upper and lower 95% confidence limit. Letters indicate significant differences based on Tukey-corrected post-hoc tests.

**Table 3.10.** The result of the ANOVA test to analyse the body size in relation to the tested temperature, FLX and 3,4-DCA conditions.

| <u>Condition</u> | <u>$\chi^2$</u> | <u>p-value</u> |
| --- | --- | --- |
| FLX | 0.34 | 0.7184 |
| 3,4-DCA | 0.00 | 0.9973 |
| Temperature | 0.01 | 0.9448 |
| Sex | 350.50 | < **0.001** |
| FLX:3,4-DCA | 0.11 | 0.8351 |
| FLX:Temperature | 0.37 | 0.7064 |
| 3,4-DCA:Temperature | 1.01 | 0.5379 |
| FLX:Sex | 0.00 | 0.9876 |
| 3,4-DCA:Sex | 1.96 | 0.3909 |
| Temperature:Sex | 2.44 | 0.3386 |
| FLX:3,4-DCA:Temperature | 4.17 | 0.2124 |
| FLX:3,4-DCA:Sex | 0.98 | 0.5434 |
| FLX:Temperature:Sex | 1.93 | 0.3939 |
| 3,4-DCA:Temperature:Sex | 0.15 | 0.8122 |
| FLX:3,4-DCA:Temperature:Sex | 0.15 | 0.8103 |

### 3.5. Fecundity

FLX exposure impacted average fecundity of female fish (Table 3.11), both in isolation and in combination with 3,4-DCA.

**Table 3.11.** The result of the mixed model ANOVA test to analyse the fecundity in relation to the tested temperature, FLX and 3,4-DCA conditions.

| Condition | $\chi^2$ | *p*-value |
| --- | --- | --- |
| FLX | 4.7548 | 0.02922 |
| 3,4-DCA | 2.7475 | 0.09741 |
| Temperature | 3.8109 | 0.05092 |
| Body Size | 0.0938 | 0.75942 |
| FLX:3,4-DCA | 4.7134 | **0.02993** |
| FLX:Temperature | 0.2727 | 0.60150 |
| 3,4-DCA:Temperature | 0.4772 | 0.48971 |
| FLX:3,4-DCA:Temperature | 0.5247 | 0.46883 |

## 4. Discussion

Animals in nature are confronted simultaneously with multiple stressors of both natural and anthropogenic origin. The overall goal of this study was to assess the sensitivity of fish to climate change, associated temperature stress, in combination with both classic pollutants (pesticides) and emerging contaminants (pharmaceuticals). Specifically, we exposed individuals of the model fish species *N. furzeri* to combinations of temperature fluctuation, 3,4-DCA and fluoxetine. Our results indicate that both traditional endpoints such as body size and fecundity as well as behavioural endpoints are affected by daily temperature variation and by the tested pollutants. The fact that certain effects were much more prominent when pollutants were combined illustrates the relevance of stressor combination studies. While the tested concentrations of pollutants did not cause excess mortality, they did induce significant behavioural changes which may carry serious fitness consequences. Therefore, our study provides an effective illustration of the need for including behavioural endpoints in ecotoxicological testing. Moreover, effects of pesticide-exposure on fish feeding behaviour in constant temperature conditions could not be confirmed under more realistic fluctuating temperature scenarios, suggesting that climate change-associated changes in temperature fluctuation should be considered to ensure the environmental relevance of ecotoxicological assessment.

### 4.1. Exposure to sublethal 3,4-DCA and FLX concentrations affects feeding behaviour in fish

Previous studies have demonstrated an anorexigenic effect of FLX in for instance fish and amphibians (Mennigen et al., 2011; Thoré et al., 2019) and this reduction in food-intake has been associated with reduced growth (Conners et al., 2009). In constrast, however, FLX exposed fish initiated feeding much faster than untreated fish in the current study. Because foraging individuals might be more conspicuous to predators (Spitz et al., 2012), we can reasonably assume that feeding is a risk-prone activity. Therefore, we hypothesise that the anxiolytic properties of FLX (Healthline, 2020) reduced stress of exposed fish and increased the tendency for risk-prone behavior (i.e. higher boldness), which may explain why exposed fish started feeding faster than unexposed individuals.

This hypothesis is consistent with personal observations which corroborate the more relaxed behaviour of FLX exposed individuals. Similar observations have been made for other aquatic organisms. Nevertheless, FLX exposure did not positively affect classic measures of fish boldness in the current study (see below), suggesting that FLX-induced anxiolytic responses alone are not sufficient to explain the observed increase in feeding behaviour upon FLX exposure under temperature fluctuation regime.

Exposure to 3,4-DCA negatively affected feeding behaviour, as fish exposed to 3,4-DCA were slower to initiate feeding compared to un-exposed fish. A parsimonious explanation could be that stress and detoxification associated with 3,4-DCA exposure may impede normal feeding behaviour (Allinson, G., & Morita, M., 1995a). However, this effect was temperature dependent and only emerged in the control temperature condition with 3,4-DCA exposed fish taking longer to initiate feeding than the un-exposed fish. Several non-mutually exclusive explanations may underpin this pattern.

Although speculative, 3,4-DCA might degrade faster under the fluctuating temperature regime due to the high peak temperatures. In support of this, Op de Beeck et al. (2017) showed that degradation-rate of the insecticide chlorpyrifos was positively correlated with ambient temperature levels., Whether this also holds under fluctuating temperature scenarios needs to be verified via water sample analyse. Alternatively, fish in the fluctuating temperature may have an upregulated metabolic rate since they experience higher maximum temperatures during certain parts of the day than fish in the control condition and this could result in an increased metabolic demand for food that counteracts the inhibitory effect of 3,4-DCA on fish feeding behaviour. After all, as ectothermic organisms their metabolism is strongly dependent on prevailing environmental temperature conditions (Brodersen et al., 2011). A similar pattern has been observed in a study of predation risk in neotropical fish (*Poecilia vivipara*) under high temperature as well (Sommer-Trembo et al., 2017).

Experiencing fluctuating temperature conditions may approximate more natural conditions for *Nothobranchius* killifish. In their temporary pool habitat, the fish experience daily temperature fluctuations of up to 15°C (T. Pinceel, unpublished data). Therefore, the fish may be more fit and able to counter the 3,4-DCA treatment more easily under fluctuating temperatures which, in turn, resulted in more alert feeding behaviour.

Importantly, the finding that 3,4-DCA exposure on fish feeding behaviour in constant temperature conditions could not be confirmed under more realistic fluctuating temperature scenarios suggests that climate change-associated changes in temperature fluctuation should be taken into account to ensure the environmental relevance of ecotoxicological assessment. To date, classic ecotoxicological tests are conducted under constant temperature conditions to ensure a high standardisation and reproducibility of results (Holmstrup et al., 2010). Doing so, however, comes at the cost of ecological relevance and may lead to an incorrect estimation of the toxicity of chemicals. The relevance of accounting for changes in temperature is further exacerbated by the projected temperature-changes under several climate-change scenarios (Klaminder et al., 2014).

### 4.2. Exposure to sublethal FLX concentrations and the relation between 3,4-DCA and body size affects activity

Also, activity was impacted by exposure to 3,4-DCA and FLX but the effects were partly dependent on body size differences among the treatments. The 3,4-DCA exposed fish overall travelled the largest distances compared to fish that were not exposed to the pesticide. This result is surprising, given that pesticide-exposure is generally reported to inhibit locomotor activity as a general indicator of stress in fish. For instance, Scheil et al. (2009) showed an impaired locomotor activity in early life stages of zebrafish upon exposure to 3,4-DCA. Although speculative, we hypothesise that the observed increase in activity upon 3,4-DCA exposure might be a hormonic effect as a widely observed yet debated, (Kaiser, 2003) phenomenon in ecotoxicology characterized by a low-dose stimulation and could be interpreted as a coping mechanism to counteract low levels of environmental stress (Calabrese & Mattson, 2017).

Total time spent for cumulative movement was greatly influenced by FLX exposure and exposed fish were more active than un-exposed fish. This seems to be a general effect that are exposed to neuroactive chemicals and is for instance consistent with the results of a study done by Ansai et al. (2016) in which chronic exposure of medaka (*Oryzias latipes*) to FLX stimulated locomotor activity. For instance, Hazelton et al. (2014), observed that FLX exposed Mussels (*Lampsilis fasciola*) (22.3 µg/L) displayed more erratic but robust and active movement in observation trials.

Maximum acceleration was unaffected by any exposure regime. Mean velocity was strongly influenced by the FLX exposure with FLX-exposed fish swimming faster than un-exposed fish. This finding is consistent with the results of a study on medaka (Ansai et al., 2016). Bossus et al. (2013) also confirmed that an amphipod species (*Echinogammarus marinus*) under FLX exposure showed a higher mean velocity.

### 4.3. Exposure to temperature variation and sublethal FLX concentrations affects boldness

Boldness was recorded and analysed through activity in the centre zone (a proxy for exposure risk to a predator attack) and latency time to enter the centre of a test aquarium. Unexpectedly, exposure to temperature variation played an important role for explaining boldness with fish from the fluctuating temperature conditions being more reserved compared to fish in constant temperature conditions. When reared under fluctuating temperatures fish were slower to enter the centre zone and spent less time in the centre zone compared to fish that were exposed to control temperatures. Potentially, fish from the temperature variation regime were forced to be accustomed to the consistent stress of fluctuating temperature (Malagié et al., 1995) and this traded off with a lower tendency for risk-prone behaviour.

Similar with the fish group from fluctuating temperature conditions, FLX-exposed fish were less prone to expose themselves in the centre of the arena than the un-exposed fish. Given the anxiolytic properties of FLX and based on previous tests with killifish (Healthline, 2020), this result was anticipated (Thoré et al., 2018). When chronically exposed to FLX, killifish displayed relatively higher anxiety towards a predation threat and they were more prone to adopt risk avoidance behaviour (Thoré et al., 2019).

### 4.4. Sex determines body size and external stressors cancel each other

Body size differed between males and females, with males being larger than females and did not differ between fish from the different pollutant and temperature treatments. Both in males and females body size is highly important for determining fitness. For males, a larger size provides benefits for coercing females into mating and for fighting other males for female attention (Cellerino et al., 2016). In *Nothobranchius* females, size is known to impact the number of eggs that can be produced daily basis (Thoré et al., 2020).

Surprisingly, we did not uncover any effects of the tested temperature regimes nor of the pollutants on body size. For both FLX and 3,4-DCA effects on growth have been demonstrated in previous studies. For instance, obstructed growth in relation to FLX exposure has been demonstrated for killifish by Mennigen and colleagues in 2011. However, they exposed fish to a much higher concentrations of 0.54 μg/L and 54 μg/L which may have caused stronger effects than our study (0.5 μg/L). Also, for 3,4-DCA effects on growth in killifish have been described in previous studies. Philippe et al. (2019), for instance, found that after exposure to 50 µg/L and 100 µg/L 3,4-DCA for 15 weeks, fish were on average 4.8% and 5.8%, respectively, larger than unexposed fish. We adopted concentrations of 50 µg/L but did not show significant difference in body size. This difference could come from lower temperature regime set by Philippe and colleagues as their temperature regime, in both variation and control, was 2°C lower (28°C was the highest temperature) than that in our study. The effects of FLX and 3,4-DCA become weaker by degradation (Op de Beeck et al., 2017; Thoré et al., 2019) when temperature is above 30°C (Op de Beeck et al., 2017). In our temperature variation setting, 30°C lasted for 3 hours during the peak time of the diurnal cycle. This high temperature effect could make both FLX and 3,4-DCA weaker due to a temperature-dependent increase in compound degradation, resulting in no strong effect from either stressor in our study.

### 4.5. Exposure to sublethal FLX concentrations affects fecundity

Fecundity was strongly impacted by FLX and exposed females produced on average less than half the eggs of unexposed fish. This result is consistent with the findings of Forsatkar et al. (2014) who tested the impact of FLX exposure on reproduction in fighting fish (*Betta splendens*). They exposed fish for seven days to a range of FLX concentration (0,54µg/L - 54µg/L). Chronic FLX exposure is also known for impair reproductive success of medaka (Brooks et al., 2003) and this is probably due to the features of SSRI that negatively alter appetite and reproduction by hormonic alteration in neurotransmitter (Malagié et al., 1995; Altamura et al., 1994; Healthline, 2020).

Surprisingly, these results are in sharp contrast to previous studies on N. furzeri, in which lifelong exposure to 0.5µg - 5 µg/L fluoxetine resulted in a doubled reproductive output (Thoré et al., 2020). These results were corroborated by several recent studies. To date, the effects of fluoxetine exposure on fish behaviour seem largely variable (Sumpter, 2014), including effects on reproductive traits (Thoré et al., 2020). Besides differences in species sensitivity and species-specificity in response to neurochemical exposure, methodological differences (including exposure dosage, duration, presence of conspecifics) might underlie this divergence.

In addition, fecundity was also impacted by 3,4-DCA exposure that disrupts reproduction (Preuss et al., 2010). When jointly exposed to FLX and 3,4-DCA females produced less than one third of eggs of un-exposed female fish. Preuss et al. (2010) found out that there was a reduction in egg production from the 3,4-DCA exposed *D. magna* and an increase in the number of aborted eggs. Hence, FLX exposure was the main reason for a decrease in egg production and 3,4-DCA amplified the effect of FLX.

## 5. Conclusion

Organisms may jointly be confronted with a multitude of stressors including climate change, pesticides and pharmaceuticals across significant portions of their life cycle in their natural environment (Cáceres, 2012; Windisch et al. 2014; Baste et al., 2012). However, typically stressor effects on organisms have been explored in isolation and at short time scales (Holmstrup et al., 2010). This leads to highly unrealistic vulnerability estimates (Fend, 2001). To obtain more realistic estimates, we need to move forward to exploring stressor interactions and combined effects of multi-stressors at long scales with practical model organism. With our study, we have taken important steps towards such added realism. We combined multiple potential stressors (temperature, pharmaceuticals and pesticides) with which organisms may be confronted in their natural environment. We also performed the study across the whole life cycle. This is important since different life stages may have different sensitivities to the same stressors, and these can only be properly assessed in whole life cycle studies. Overall, with our study we demonstrate that the killifish *N. furzeri* can be a time and cost-efficient model for realistic whole life cycle and multi-generational studies.

Some of our findings, especially relevant to feeding behaviour and body size in relation to pollutant exposure, were not consistent with our expectations based on the available literature (Philippe et al. 2018; Thoré et al. 2019). Inconsistency of FLX effects on aquatic organisms across studies and test species has been recognised for some years now (Thoré et al., 2020). Likely this at least partly derives from a lack in test standardisation. Therefore, ecotoxicology, especially regarding long term testing on fish, needs a reboot in the form of standardised culturing protocols for model organisms and adapted OECD guidelines for long term and multi-generational testing. Also, our results showcase that while classic ecotoxicological endpoints such as mortality and growth may sometimes not be affected by low and ecologically relevant pollutant concentrations, other sublethal endpoints which also carry fitness effects such as behavioural expression may still be affected. Therefore, also such endpoints should become more embedded in standard ecotoxicological testing.

Unfortunately, we had to terminate our experiment after a single generation. However, an interesting next step would be to now also track egg survival up to the next generation and hatching success and survival/life history across the next generation. After all, we uncovered significant effects on fecundity of the different tested stressors and even in eggs that were produced other effects such as reduced survival or hatching success may yet emerge. Still, even the findings of our time-limited study are relevant and valuable in several ways. Combined, they demonstrate the need to include tracking of behavioural endpoints in whole life stressor combination studies.

## Supporting information

Body size

Fecundity

Fecundity and body size

Feeding

Growth

Open field

Open field and feeding

## List of abbreviations

3,4-DCA: 3,4-dichloroaniline
ACR: Acute to chronic ratio
EC50: Effective concentration 50 percent of all individuals
FLX: Fluoxetine
IPCC: Intergovernmental Panel on Climate Change
LC50: Lethal concentration for 50 percent of all individuals
LMM: Linear mixed model
LOEC: Lowest observable effect concentration
NOEC: No observed effect concentration
SSRI: Selective serotonin reuptake inhibitor

## Acknowledgements

This manuscript was written as a master’s thesis (academic year 2019-2020), part of MSc Sustainable Development under the supervision of Professor Luc Brendonck, Dr Tom Pinceel and Dr Eli Thoré at KU Leuven, Belgium. I express my gratitude to my supervisors, technicians and colleagues at Kolenmuseum, Department of Biology, KU Leuven.

---

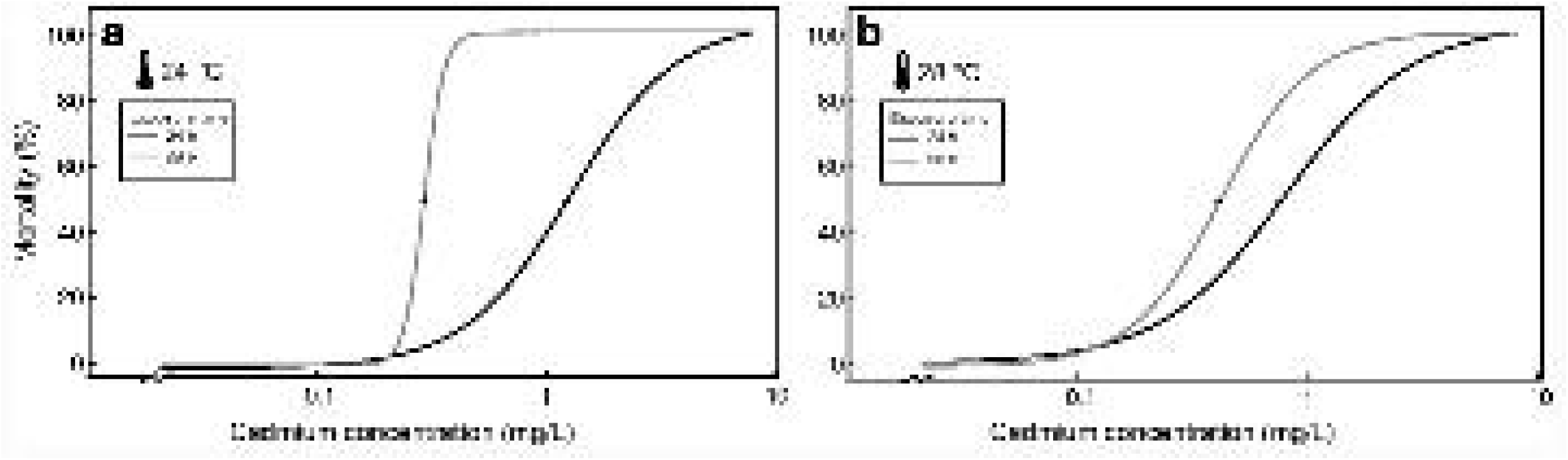

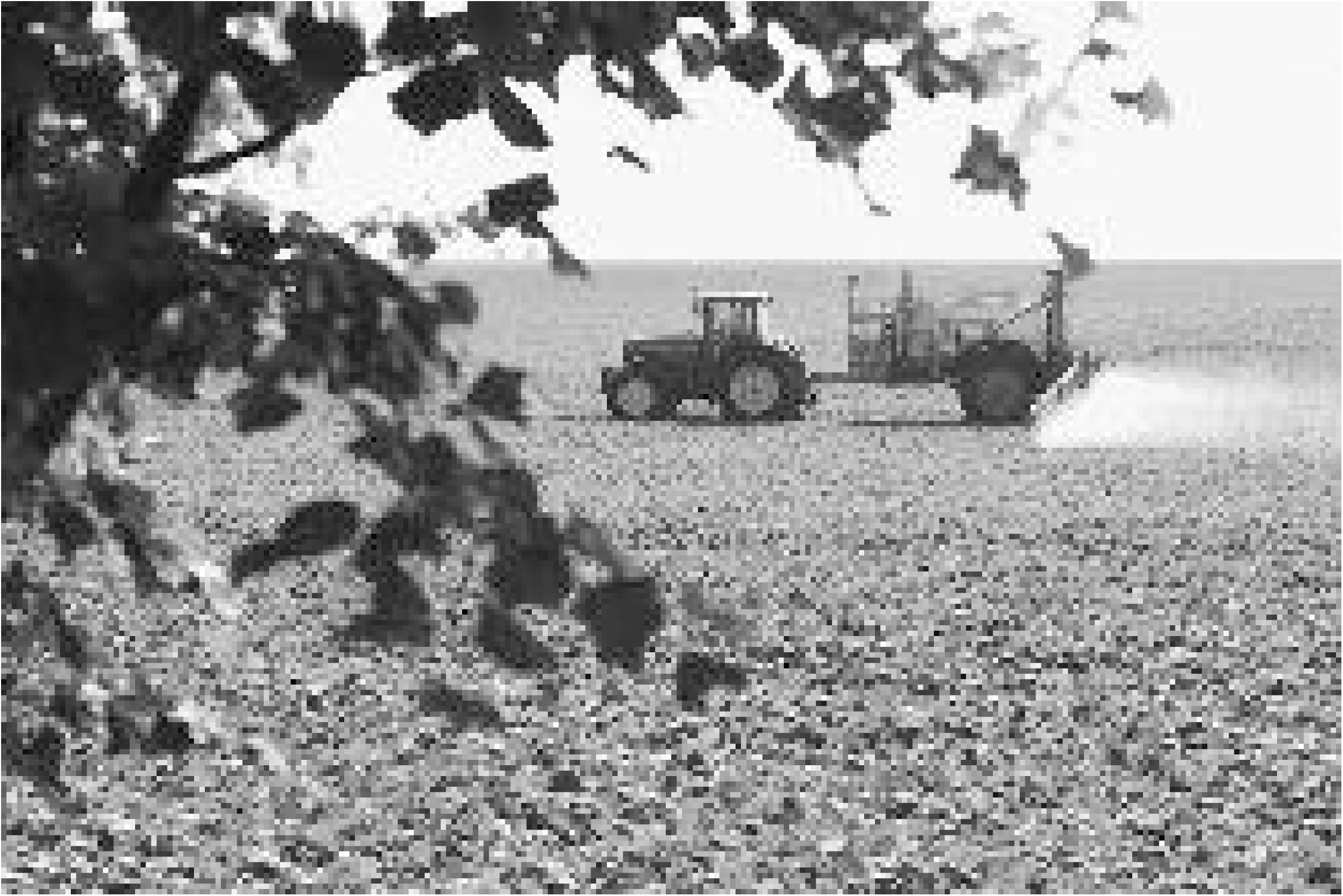

## Notes

### Competing Interest Statement

The authors have declared no competing interest.

### Summary of Updates

The section within the institute changed whereas the institutional affiliation remains the same.

